# The structure of Lepidoptera-plant interaction networks across clades, life stages, and environmental gradients

**DOI:** 10.1101/2022.11.10.516059

**Authors:** Hsi-Cheng Ho, Florian Altermatt

## Abstract

**Aim:** Integrate biogeographic and ecological knowledge to understand the spatial-structural patterns of plant-insect interaction networks at the landscape scale.

**Location:** The 36,000 km^2^ German state of Baden-Württemberg, Central Europe.

**Methods:** We integrated extensive data of Lepidoptera-plant occurrences and interactions to inferentially construct local interaction networks across Baden-Württemberg, considering in total 3148 plant and 980 Lepidoptera species, covering butterflies, Noctuid moths, Geometrid moths and Bombycoid moths. We quantified clade- and life-stage-specific network structures and related these features to GIS-informed environmental conditions, thereby revealing the spatial (environmental) patterns and potential drivers of networks’ structural variation across the landscape.

**Results:** Spanning the same environmental gradients, Lepidoptera clades and life stages can form various interaction structures with food plants and exhibit distinct spatial-structural patterns. For all major Lepidopteran groups, except Geometrid moths, potential diet across life stages tended to broaden toward low-elevation farmlands. The larval and adult networks of butterflies became less modular with farmland coverage; the same for adult Noctuid moths, but the inverse for adult Geometrid moths. With increasing elevation, the larval and adult networks of Noctuid moths became less and more modular, respectively, whereas Geometrid adult networks became more modular. While the adult dietary niche of butterflies was more overlapped at low elevation, those of Noctuid and Geometrid moths further associated with land cover and were more overlapped toward low- and high-elevation farmlands, respectively.

**Main conclusions:** Environmental factors and biotic interactions together shape ecological communities. By particularly accounting for species-interaction contexts, we revealed the spatial-structural patterns of Lepidoptera-plant networks along geo-climate and land-cover gradients, where the shaping mechanisms likely include both evolutionary (e.g., resource-consumer co-evolution) and ecological (e.g., competitive exclusion) processes and are specific to Lepidoptera’s clade or life stage. Such biogeographical structural patterns provide ecological and conservation implications at both species and community levels, and can indicate the potential response of Lepidoptera-plant communities to environmental changes.

## Introduction

Species occurring in the same habitat or site can interact inextricably with each other, forming interaction networks whose features affect species persistence as well as stability and functioning of the whole community (Bascompte, 2009; Thompson et al., 2012). For a needed better management of biodiversity and ecosystem services in the Anthropocene, it is essential to understand these interaction structures and their dependencies on biotic and abitotic factors across realistic landscape scales (Thompson et al., 2012; Tylianakis et al., 2010).

Of particular interest are plant-insect interaction networks because they are well-studied, and cover many species who often play important roles in ecosystems. Of all plant-insect interactions, Lepidoptera-plant system are arguably best-studied, and their understanding can likely be projected across many further plant-insect interaction networks. Lepidoptera form a hyper-diverse order and are a prevailing component in terrestrial ecosystems worldwide (Scoble, 1995). The holometabolous Lepidopterans play “dual roles” interacting with plants. Specifically, in most cases, their larvae are herbivorous and some are powerful pests, whereas most adults utilise floral resources and serve as key pollinators (Boggs, 1987). This covers both antagonistic and mutualistic relationships and often across the same Lepidoptera-plant species pairs (Altermatt & Pearse, 2011). Thus, disentangling the potentially divergent network patterns between Lepidoptera life stages interacting with the same local plant assemblages could have major eco-functional implications (Astegiano et al., 2017). Lastly, Lepidopterans and their food plants have co-evolved into strong mutual dependencies (Pearse & Altermatt, 2013a; Weiblen et al., 2006). Their interactions are therefore crucial to both sides, making the Lepidoptera-plant system somewhat operate distinctively, and suitable to be viewed as a standalone unit. Strong dependencies also imply high potential of cascading effects. Hence, the system should be especially prone to (and thus appropriate for studying) environmental influence cascading across the whole interaction network (Pearse & Altermatt, 2013b).

Past studies on plant-insect networks suggest a typical modular structure, reflecting that sets of insects tend to interact with respective sets of plants but rarely outside such modules that they formed (Astegiano et al., 2017; Braga et al., 2018; Olesen et al. 2007), and such pattern is echoed in individual Lepidoptera-plant network studies (López-Carretero et al., 2014). However, these studies are mostly conducted on a local scale and cannot capture the full spatial or environmental reliance of network structure (Pellissier et al., 2018; Tylianakis & Morris, 2017). Meanwhile, there have been large-scale studies investigating the geographical distributions of Lepidopterans and their food plants, often as global or continental integrations (e.g., Carvajal Acosta & Mooney, 2021). Yet, these studies usually focus on a few species’ co-occurrence and thus do not inform network understandings (but see Narango et al., 2020). While it is often the environmental gradients spanning between local and continental scales that shape species distributions within a landscape (Hanspach et al., 2014; Jones, 2011), the corresponding interaction-network realisation remains largely unexplored.

Here, we integrate biogeographic (i.e., who occurs where) and ecological (i.e., who interacts with whom and how) knowledge to examine Lepidoptera-plant network structure at a landscape scale. Specifically, we target the geographic variation of network structure in three aspects: (1) between Lepidoptera life stages, (2) among Lepidoptera clades, and (3) along selected environmental gradients. We thereby disclose the structure of antagonistic and mutualistic interactions formed by the same local assemblages of Lepidoptera and plants, as well as the structural differences among Lepidoptera clades. More importantly, we identify which and how key environmental drivers, such as geo-climate and land cover, known to influence biodiversity (Mantyka-Pringle et al., 2015), may have shaped the composition and interaction structure of Lepidoptera-plant communities through life-stage- or clade-specific traits or processes.

We based our analysis on extensive, long-term, and virtually complete empirical datasets on the occurrences and interactions of Lepidoptera and plants in Baden-Württemberg, Germany (Central Europe). We applied the metaweb approach (see *Methods*) to integrate occurrence and interaction information to construct local Lepidoptera-plant networks of each Lepidoptera clade and each life stage, on a grid basis across the study area. Environmental information of the same grid cells was derived from geographic information systems (GIS). We quantified the structure of local networks with a selected set of frequently studied network metrics (see *Methods*) and identified different structural features across Lepidoptera clades and life stages. Finally, we related the detected structures to geo-climate and land-cover gradients to reveal the environmental reliance of the spatial-structural patterns of Lepidoptera-plant networks.

## Material and Methods

### Study area and its environment

Our study area is the German state of Baden-Württemberg in Central Europe, spanning 35,752 km^2^ and a vertical range from 85 to 1,493 m above sea level. The study area was spatially resolved to 10’ longitude × 6’ latitude grids (roughly 10 × 10 km^2^). Hereinafter, we refer to such grid cells as the “local” scale, which presents the resolution of our species occurrence, environmental variables, and constructed interaction networks (see further below). In total, our data covered 310 such grids (Fig. 1).

**Figure 1.**
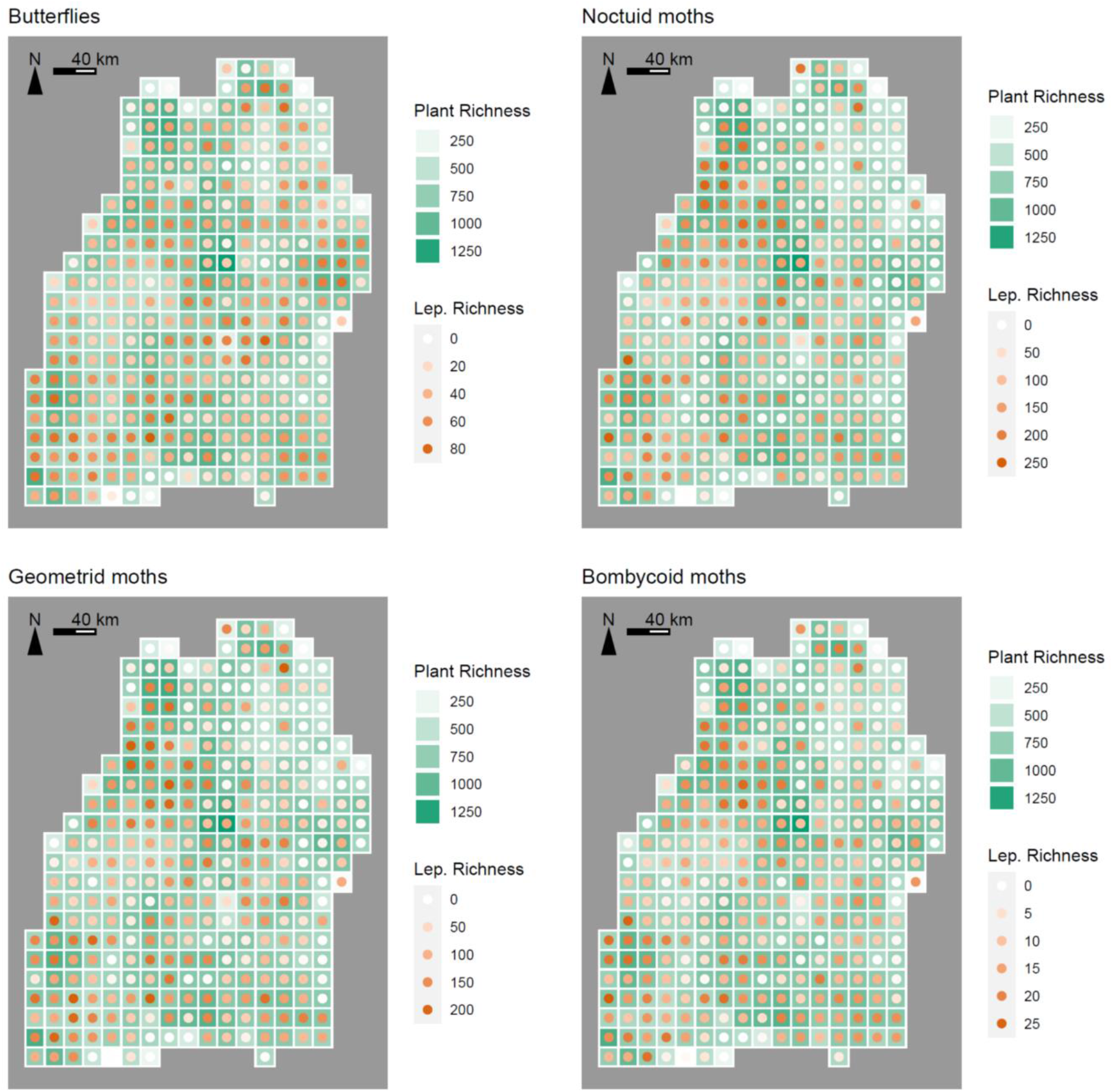
The richness of plants and Lepidoptera recorded in each of the 310 grid cells across Baden-Württemberg. The four focal clades of Lepidoptera, namely butterflies, Noctuid moths, Geometrid moths, and Bombycoid moths, are each presented in a subfigure. The spatial richness patterns of the three moth clades were highly consistent despite their various ranges of richness, while butterflies exhibited a somewhat different pattern. For their correlations, see Fig. S3.

We extracted geographic, climate, and land-cover information of the study area from available GIS databases, then spatially resolved the information to match the same grids abovementioned. The variables included: mean elevation of each grid cell from *Shuttle Radar Topography Mission* (SRTM, NASA/NGA), mean temperature of each grid cell from *CHELSA*, and local land cover from *CORINE Land Cover* (CLC, EEA), all averaged over the years of 2005–2015. For land cover, we dropped cover types that generally occupy only a minimal area per grid cell, thereby focusing on three major cover types in subsequent analyses: forests (mean coverage 37.2 %, range 1.3–88.7 %), farmlands (49.6 %, 5.4–93.9 %), and urban areas (10.4 %, 0.2–59.3 %).

### Data on Lepidoptera, plants, and their interactions

Our study considers Lepidoptera (butterflies and moths) and plant species that were recorded in the study area within a time window of three decades (1985–2014). These occurrence data were derived from respective long-term monitoring and natural history surveys encompassing all the local grid cells (Database *Arbeitsgruppe Schmetterlinge Baden-Württembergs am SMNK* https://www.schmetterlinge-bw.de for Lepidoptera; *Bundesamt für Naturschutz 2010 & Floraweb* for native and naturalised plants). We note that, the sampling followed a haphazard sampling regime, Lepidoptera monitoring was so intensive that grids were generally well-sampled for all taxonomic groups considered. Thus, although our local scale and corresponding environmental information spatially resolve the finest to the size of grids, the recorded Lepidoptera species therein should reflect local (*sensu stricto*) assemblages shaped by localised processes, yet integrated at the sampling grid. Also notably, by compiling present-absent occurrence information across such a time window, we dropped ancient records where the species may no longer exist, and took advantage of the more systematically and homogenously sampled data from modern era (Supplementary Information Fig. S1). Meanwhile, we still look at a long enough timeframe to capture the occurrence of rare species that possibly only detected intermittently. In total, the dataset contains local occurrences of 980 Lepidoptera species and 3148 plant species (a few as aggregated species-complex). We then focused on four main clades of Lepidoptera in Baden-Württemberg, namely butterflies (Papilionoidea, incl. Hesperiidae, Lycaenidae, Nymphalidae, Papilionidae, and Pieridae), Noctuid moths (Noctuoidea, incl. Arctiidae, Lymantriidae, Noctuidae, Nolidae, Notodontidae, and Pantheidae), Geometrid moths (Geometroidea), and Bombycoid moths *sensu lato* (Bombycoidea and Lasiocampoidea, incl. Endromidae, Lasiocampidae, Lemoniidae, Saturniidae, and Sphingidae). These clades form monophyletic groups (Kawahara et al., 2019) and are relatively diverse in the area, making them suitable subjects for among-clade comparisons. We interpreted a species’ record as an occurrence of its both life stages (larval and adult, respectively) in a given grid cell. Consequently, we expected some sampling bias towards butterflies due to their diel-activity patterns.

To construct the Lepidoptera-plant interaction network, we compiled the dietary information of Lepidopterans mainly based on the work of Ebert (1991–2005) and further complemented with other references and personal observations (Altermatt & Pearse, 2011; Pearse & Altermatt, 2013a, 2013b). This covered both their larval usage of host plants and adult usage of floral resources. Such dietary information was grounded on empirical observations of feeding under natural, un-manipulated field situations, recorded by professional entomologists over a course longer than 50 years and with more than 2.3 million observations of individual Lepidoptera-host interactions (Ebert, 1991–2005; Pearse & Altermatt, 2015). We converted the recorded interactions into binary (i.e., interact or not) dietary matrices between Lepidoptera and plants, larval and adult stages separated. Given the enormous sampling intensity, and since we extracted only the binary information from accumulative records, we treated our dietary matrices as being complete and free from under-sampling (Pearse & Altermatt, 2015).

We subsequently applied the “metaweb” approach (Ho et al., 2022; Saravia et al., 2022) to extrapolate local co-occurrences of Lepidoptera and plants to local interaction networks, taking our dietary matrices as the respective larval and adult metawebs. The approach assumes that a feeding interaction indicated in the metaweb will realise if the corresponding Lepidoptera-plant species pair co-occur locally, that is in the given grid cells looked at. In other words, every Lepidoptera species at a given life stage has a fixed set of potential regional food plants, which could be used locally if co-occurring. This assumption embraces the concept that interactions are driven by matching functional characteristics (e.g., chemical tolerance in Lepidoptera, see Després et al., 2007), and collapses potential intraspecific variations of these characteristics at the species level (Ho et al., 2022). In our case, we assumed no spatial structure and mobility restrictions of the species within the local grid cells, that is, assumed that all species present can interact. We inferred the realisation of interactions between locally (grid-level) co-occurring Lepidoptera and plants using the metawebs, thereby constructing local Lepidoptera-plant networks for each Lepidoptera clade and life stage. Species without local interacting partners (e.g., Lepidoptera adults without functioning proboscis, or plants that are not food to any focal Lepidoptera) were excluded from the networks. In other words, although at each local site there was a fixed plant assemblage, for each focal clade and life stage only the respective food-plant subset was accounted. Our constructed local networks essentially reflect potential (not locally-sampled) interactions, but importantly within empirically-derived boundaries of realistic interactions and co-occurrences. Thus, beyond the local scale, they can guide an unbiased exploration of spatial-structural patterns of networks resulting from compositional difference of local communities.

### Spatial diversity and network analyses

Based on our occurrence data, we derived species diversity (richness) of plants and each focal Lepidoptera clades across all 310 local grid cells. We analysed the correlations among these grid-wise richness to check if they exhibited correlated spatial diversity patterns. Such patterns were also visualised on regional grid maps of diversities (Fig. 1). Then, to disentangle environmental drivers that may have contributed to these spatial diversity patterns, we conducted a principal component analysis (PCA) with grid-wise mean elevation, mean temperature, and proportional land cover of forests, farmlands, and urban areas as the explaining variables (N = 310). The identified PC1 and PC2 (Fig. 2; from a retrospect, respectively reflected geo-climate and land-cover gradients) were then taken as environmental predictors for a series of general linear model (GLM) analyses on Lepidopterans’ and plants’ richness. A significant non-zero slope detected in the analyses would indicate the corresponding PC’s significant influence. The richness × PC interaction terms were included in the Lepidopteran analyses to examine potential slope difference among Lepidoptera clades.

**Figure 2.**
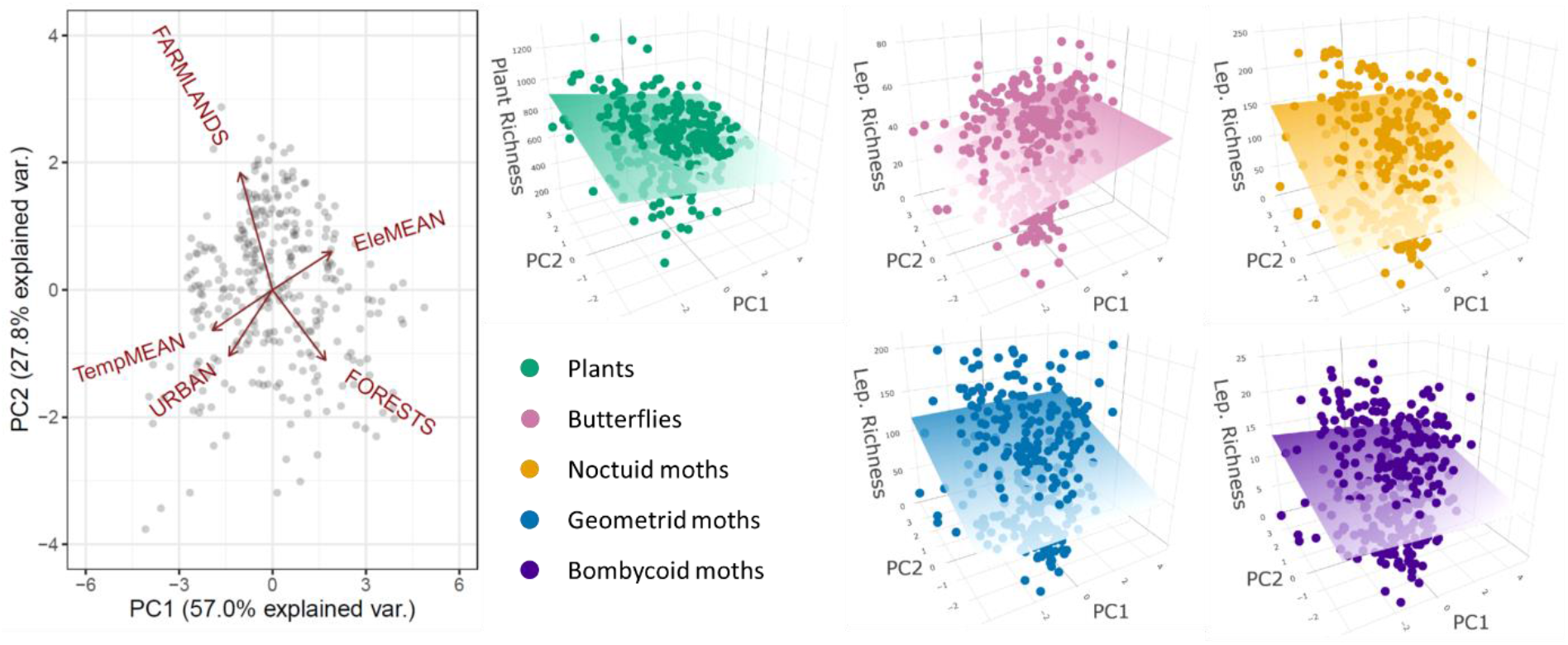
The first two principle components of environmental factors (PC1 and PC2, considered as axes of gradients), and the richness of plants and the four Lepidoptera clades across these two axes as 3D scatterplots. The planes in the 3D scatterplots are regression planes of the observed values, whose colour fade toward the low-value end. Corresponding stats with separated regressions against each PC are provided in Fig. S7 & S8.

With our constructed local Lepidoptera-plant networks, we quantified their structure with a selected set of structural metrics: mean generality of Lepidopteran consumers, network modularity, and mean dietary niche overlap of Lepidopterans. Mean generality reflects the mean diet breadth of Lepidoptera with a given local plant assemblage. Modularity indicates the prevalence of a modular structure within a network (ecological relevance see *Introduction*; calculations see Section S1). Lepidopterans’ dietary niche overlap was evaluated by their local food-plant usage (Section S1). The higher the overlap, the stronger the Lepidopterans’ diet competition. Niche overlap could also be evaluated from the plants’ perspective, indicating their apparent competition (in antagonistic larval networks) or pollinator competition (in mutualistic adult networks). We here focused on the Lepidoptera’s perspective since the values from both perspectives were essentially highly correlated (Fig. S2). We also additionally addressed network connectance and nestedness, and present the relevant content in Section S2. All these metrics are frequently examined in ecological network studies (e.g., Ho et al., 2022; Thébault & Fontaine, 2010), as they capture not only structural features but also ecological implications spanning organismal (e.g., diet breadth) and community (e.g., niche differentiation) scales.

To resolve whether there are structural differences among the networks of Lepidoptera clades, and whether environmental drivers may have shaped their spatial-structural patterns via influencing community composition, we mirrored the analyses in analogy to species richness, performing a series of GLM of network metrics among Lepidoptera clades and each against the two PCs, respectively. Notably, for the hereinafter network analyses we excluded small local networks with either plant or Lepidoptera richness fewer than ten. Small networks tend to generate unreliable and artifactual structural-metric readings (Dormann et al., 2009) that distort the interpretation. Thus, across all 310 grid cells, the number available local networks (i.e., sample size) of each Lepidoptera clade-stage combination were: 272 and 271 for larval and adult butterflies, 251 and 239 for Noctuid moths, 252 and 236 for Geometrid moths, and 146 for larval Bombycoid moths (no remaining adult network, as the adult Bombycoid moths considered largely do not have a functional proboscis).

With the detected spatial-structural patterns of local networks, to further disentangle the possible shaping mechanisms of such patterns, we applied two null models to simulate respective randomised networks. These model-generated “counterparts” were designed to be comparable to the local networks (whose metric readings were “observations”). The first null model is “re-assembled”: with a local network at a given grid cell as the input, this model randomly draws Lepidoptera and plants from the study area’s respective species pools, and uses our metawebs to wire them into a new local network with exactly the same number of species in both trophic levels as the input. Therefore, networks generated by this model are re-assembled Lepidoptera-plant communities where the interactions remain biologically realistic (i.e., based on realistic metawebs), but the real-world species-environment reliance is destroyed (i.e., local species assemblages are randomly composed instead of driven by the environment). The second null model is “shuffled”: with a local network at a given grid cell as the input, this model keeps the same number of nodes at each trophic level and the same total number of links, but rearranges the links among the nodes, thereby generating a new network. Therefore, networks generated by this model have neither biologically realistic identities nor interactions of species—both the real-world Lepidoptera-plant interdependences and species-environment reliance are destroyed. For each observed local networks, we generated twenty randomised counterparts with each null model. Looking at structural metrics’ readings, if an observation lies within the confident interval estimated from its twenty “re-assembled” counterparts, such observed structure is likely not shaped by real-world species-environment reliance, because the randomly re-assembled communities already exhibit the same structure without realistic community composition. In contrast, an observation outside such confident interval indicates a significant (positive or negative) contribution of real-world species-environment reliance on the structural feature. Similarly, an observation outside the confident interval of “shuffled” counterparts indicates a significant collective contribution of real-world species-environment reliance and Lepidoptera-plant interdependences. We made comparisons between observed networks and null-model counterparts across the five focal network metrics to examine if the observed spatial patterns were driven by these mechanisms. An illustration of how we unified and visualised such comparisons using Z-scores and 3D regression plots is given in Fig. 3.

**Figure 3.**
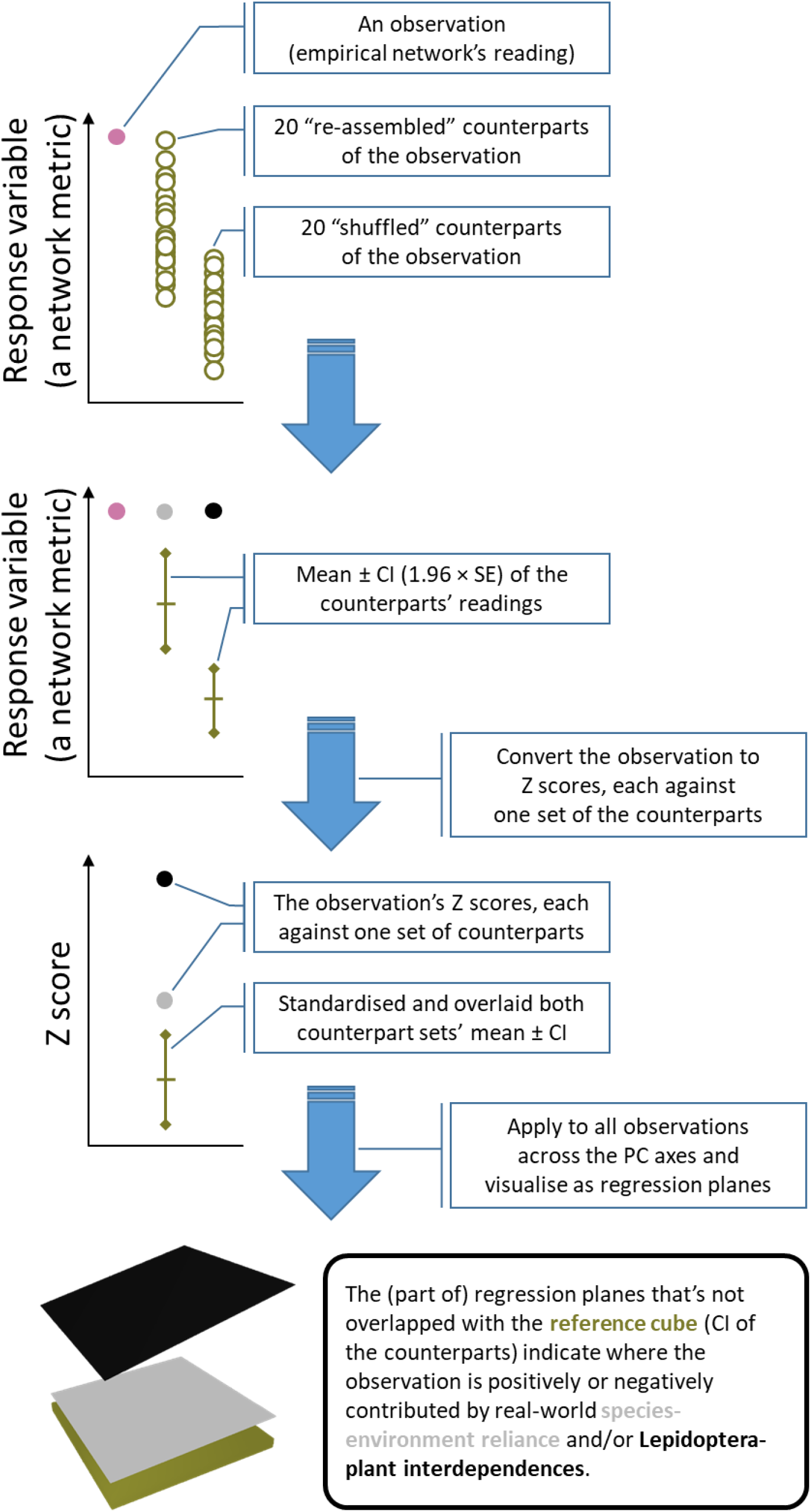
An illustration of how observations of network metrics are converted to Z scores against the two randomised counterparts (generated by “re-assembled” and “shuffled” null models, see *Methods*) to disentangle the contribution of real-world species-environment reliance and Lepidoptera-plant interdependences on shaping empirical network properties.

All quantifications, analyses, and plotting were performed using R version 4.2.1 (Core R Team, 2013). Relevant information, including applied packages and functions, is described in Section S1. The R scripts performing these tasks are accessible at the provided online repository.

## Results

Across all 310 studied grids, the local (per grid cell) richness of plants averaged 730.6 ± 181.8 species (mean ± SD), while the local richness of Lepidoptera averaged 38.1 ± 20.9 species for butterflies, 93.1 ± 69.2 for Noctuid moths, 81.9 ± 61.1 for Geometrid moths, and 10.0 ± 7.0 for Bombycoid moths (Fig. 1). For each of the four focal Lepidoptera clades, local Lepidopteran richness was significantly positively correlated with local plant richness (Fig. S3). Looking into further details, the three moth clades exhibited highly consistent spatial diversity patterns despite their various range of richness, whereas the butterfly clade somewhat differed from the moths (with ∼0.9 high correlation coefficients among moth clades, while ∼0.6 moderate coefficients between moth and butterfly clades; see Fig. S3). Fig. 1 visualises these patterns and depicts species diversity hotspots (dark orange dots) of moth clades were well-aligned with each other, but less aligned with butterflies.

The potential environmental contributors of observed spatial diversity differences among focal clades were revealed by our principle component analysis (PCA) of environmental factors. The first two PC axes together explained 84.8% of the environmental variation among local grids. The PC1 mostly captured the geo-climate variation, i.e., mean elevation and temperature, while the PC2 the land-cover variation, i.e., the coverage of forests, farmlands, or urban areas (Fig. 2). Informed by the regressions against the two PC axes, the richness of butterflies was more influenced, positively, by PC1 (R^2^ = 0.04, p < 0.001; Fig. S8), whereas the richness of the three moth clades were consistently more influenced, negatively, by PC2 (R^2^ = 0.05, 0.04, 0.02 and p < 0.001, < 0.001 and = 0.021 for Noctuid, Geometrid, and Bombycoid moths, respectively; Fig. S8). In other words, there were more butterfly species toward high-elevation and low-temperature areas (note that the whole study area, and thus the mean elevation of every grid cell, was below 1,500 m a.s.l.), while more moth species toward farmland-dominated lands (Fig. 2 & Fig. S8).

Then, with the constructed clade-specific Lepidoptera-plant local networks (sample sizes see *Methods*; Fig. S4), we quantified and associated their structural features to the environmental PC axes, and compared the observations with null models (see *Methods &* Fig. 3). With respect to the mean generality (i.e., diet breadth) of Lepidoptera in the interaction networks, larval butterflies had the narrowest diets among the four clades, but conversely with the broadest diets as adults (p < 0.001 for comparing with any moth clade; Fig. S9). The tendencies of mean generality varying along the PC axes were generally consistent across the four Lepidoptera clades at the larval stage. That is, higher ≥for PC1 and PC2 in butterflies, and R^2^ = 0.02 and p = 0.03 for PC1 in Geometrid moths; Fig. 4 & Fig. S9). At the adult stage, butterflies and Noctuid moths exhibited qualitatively similar patterns as of the larvae (R^2^ = 0.06 and 0.03, p < 0.001 and = 0.008, respectively, for PC1 and PC2 in butterflies, and R^2^ = 0.03 and p = 0.006 for PC1 in Noctuid moths; Fig. 4 & Fig. S9). However, an inversed pattern was detected in adult Geometrid moths, i.e., having broader diets toward high-elevation forests (R^2^ = 0.02, and p = 0.058 and 0.031, respectively, for PC1 and PC2; Fig. 4 & Fig. S9). The null-model analyses further showed that the larval generality patterns of the four clades were positively driven by real-world species-environment reliance (i.e., realistic species assemblages driven by the environment), and similarly for the adult pattern of butterflies. Instead, the adult patterns of Noctuid and Geometrid moths were negatively driven by such reliance (Fig. 4).

**Figure 4.**
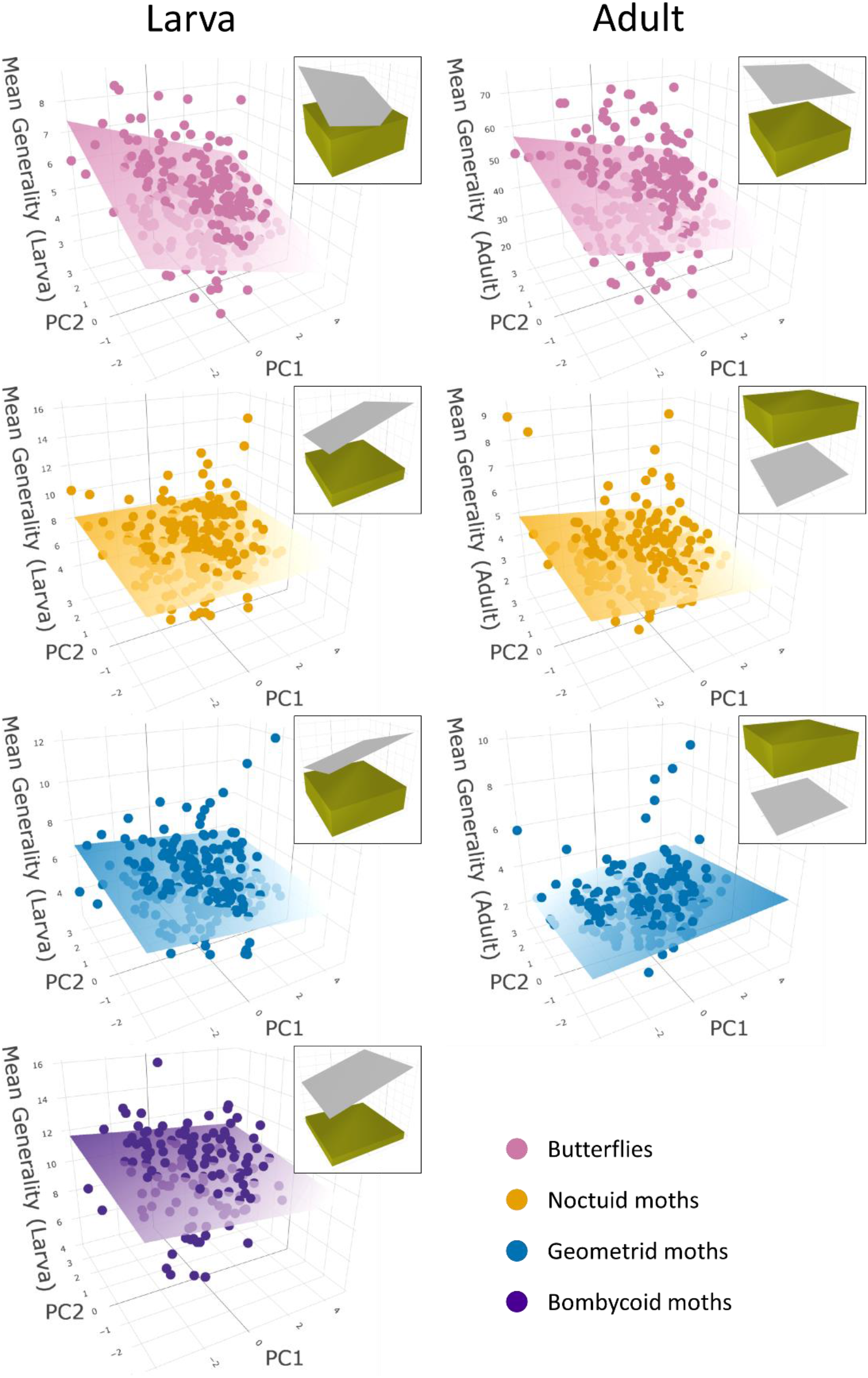
Mean generality (diet breadth) of Lepidoptera in the local networks across environmental gradients as 3D scatterplots, sharing the same environmental PC axes (as in Fig. 2) but respective metric reading axis. The planes in the 3D scatterplots are regression planes of observed values, where their colour fade toward the low-value end. Corresponding stats with separated regressions against each PC are provided in Fig. S9. Corresponding null-model analysis against randomised counterparts is given at each subplot’s top-right (details see *Methods* and Fig. 3). Note that since the total number of links, and thus mean generality, were fixed to empirical values in the “shuffled” null model, only the “re-assembled” planes appear in the null-model analyses.

Regarding network modularity, butterflies formed the most modular networks among the four clades as larvae, but the least modular ones as adults (p < 0.001 for comparing with any moth clade; Fig. S10). The modularity patterns along the PC axes were inconsistent among the clades and depending on the life stage. On the one hand, the larval network of all clades tended to become less modular toward high PC2, i.e., increasing farmland coverage, but only the response of butterflies was significant (R^2^ = 0.05, p < 0.001; Fig. 5 & Fig. S10). The modularity of both Noctuid’s and Geometrid’s larval networks were influenced by PC1 but in opposite directions, such that the former became less modular while the latter became more modular toward high PC1, i.e., high elevation and low temperature (R^2^ = 0.01, p = 0.062 in Noctuid moths and R^2^ = 0.02, p = 0.040 in Geometrid moths; Fig. 5 & Fig. S10). On the other hand, the adult networks of all clades (excl. Bombycoid moths; same for all following statements of adult-network comparisons) were influenced by PC2, such that butterflies and Noctuid moths formed less modular networks along PC2 (R^2^ = 0.08, p < 0.001 and R^2^ = 0.01, p = 0.078, respectively; Fig.5 & Fig. S10), while the inverse for Geometrid moths (R^2^ = 0.02, p = 0.020; Fig. 5 & Fig. S10). Along PC1, in contrast, only Noctuid’s adult networks became significantly more modular (R^2^ = 0.03, p = 0.004; Fig. 5 & Fig. S10). The null-model analyses showed that both real-world species-environment reliance and Lepidoptera-plant interdependences (i.e., realistic Lepidopteran diets) shaped network modularity in similar ways, which differed across clades and life stages. In general, these realistic biological constraints contributed positively to the modularity of butterflies’ larval networks and negatively to their adult networks, while the opposite in the moth clades (Fig. 5).

**Figure 5.**
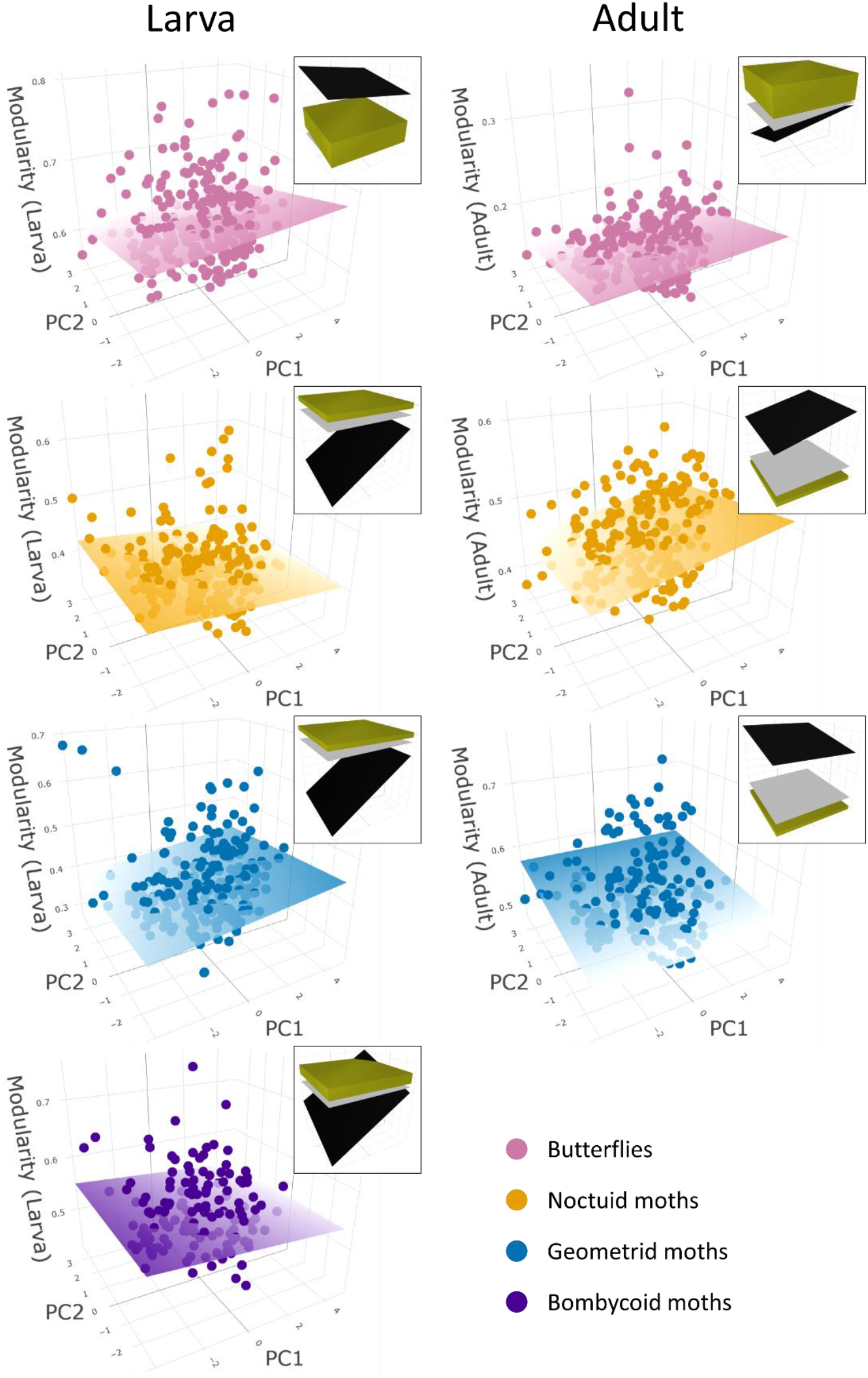
Modularity of the local networks across environmental gradients as 3D scatterplots, sharing the same environmental PC axes (as in Fig. 2) but respective metric reading axis. The planes in the 3D scatterplots are regression planes of observed values, where their colour fade toward the low-value end. Corresponding stats with separated regressions against each PC are provided in Fig. S10. Corresponding null-model analysis against randomised counterparts is given at each subplot’s top-right (details see *Methods* and Fig. 3).

As for the Lepidopterans’ dietary niche overlap, butterflies again had a relatively drastic divergence between life stages, as their larval diets were the least overlapped while the adult diets the most overlapped among all clades (p < 0.001 for comparing with any moth clade; Fig. S11). The environmental drivers were mostly not influential to niche overlap in larval networks. Only the diets of larval butterflies became significantly less overlapped toward high PC1 (R^2^ = 0.09, p < 0.001; Fig. 6 & Fig. S11). On the contrary, the PCs were more influential at the adult stage. Along increasing PC1, niche overlap increased in Geometrid moths (R^2^ = 0.02, p = 0.023) but decreased in butterflies (R^2^ = 0.01, p = 0.076) and Noctuid moths (R^2^ = 0.07, p < 0.001; Fig. 6 & Fig. S11). Along increasing PC2, diets in Noctuid and Geometrid moths’ adult networks became more overlapped (R^2^ = 0.05, p < 0.001 and R^2^ = 0.02, p = 0.056, respectively; Fig. 6 & Fig. S11). The null-model analyses again revealed consistent yet clade- and life-stage-specific contributions of both real-world species-environment reliance and Lepidoptera-plant interdependences. These constraints contributed positively to niche overlap in the larval networks of all clades, as well as in the adult networks of butterflies, while negatively in the adult networks of Geometrid moths (Fig. 6).

**Figure 6.**
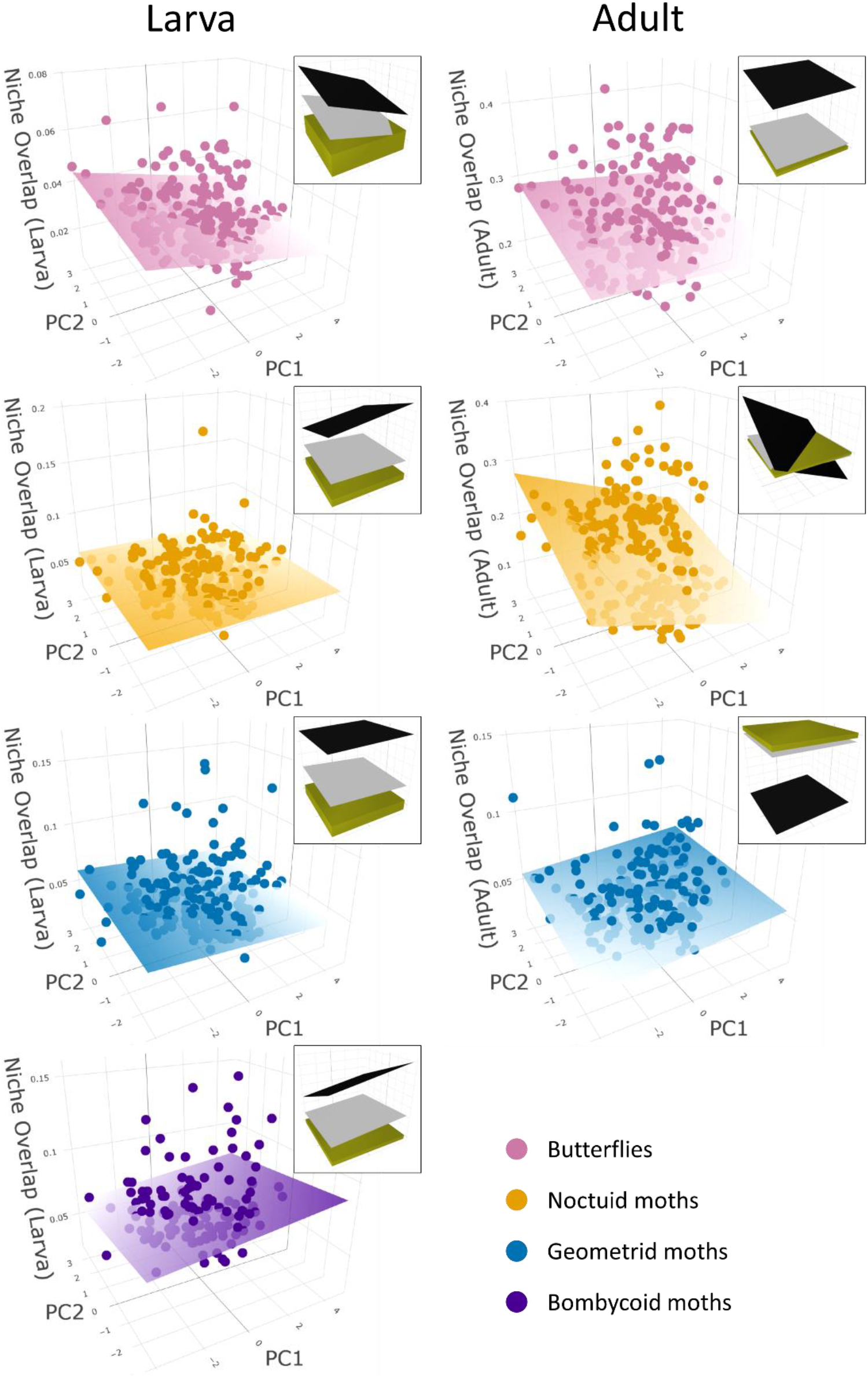
Dietary niche overlap of Lepidoptera in the local networks across environmental gradients as 3D scatterplots, sharing the same environmental PC axes (as in Fig. 2) but respective metric reading axis. The planes in the 3D scatterplots are regression planes of observed values, where their colour fade toward the low-value end. Corresponding stats with separated regressions against each PC are provided in Fig. S11. Corresponding null-model analysis against randomised counterparts is given at each subplot’s top-right (details see *Methods* and Fig. 3).

## Discussion

Based on extensive datasets of species occurrences and interactions, we brought together biogeographic and ecological knowledge to reveal landscape-scale spatial patterns of species diversity and potential interaction networks of Lepidoptera-plant communities. Specifically, these patterns reveal how abiotic (e.g., climate and land cover) and biotic (e.g., resource-consumer reliance) contexts may together determine the composition and structure of local interaction networks.

Overall, local Lepidoptera richness, regardless of clades, positively correlated with plant richness. These patterns likely reflect Lepidoptera’s strong dependence on their interacting plants rather than the reverse, given that most Lepidopterans across life stages are sole and relatively specialised plant feeders (Pearse & Altermatt, 2013a), while plants rarely rely only on Lepidoptera adults to pollinate them (Gibson et al., 2006; Memmott, 1999). Thus, more diverse local plant assemblages can potentially support the existence of more Lepidoptera species (Axmacher et al., 2009). Still, we detected systematic difference in spatial diversity patterns among Lepidoptera clades, such that butterfly richness was driven more by geo-climatic environmental factors (PC1; more diverse toward higher elevation), while the richness of the moth clades more by land covers (PC2; more diverse toward farmland-dominant lands). Our butterfly altitudinal richness pattern echoes other independent butterfly studies in nearby geographical regions and likely captures the positive richness-elevation association before reaching a mid-elevation (∼1,500 m a.s.l.) peak or plateau (see dome-shape relationships observed in Gallou et al., 2017; Ho et al., 2022). Since this pattern was butterfly-specific, it is unlikely due to a geometric Rapoport effect (Beck et al., 2016; Colwell & Hurtt, 1994) that would have prevailed also in other clades, but instead reflecting biological constraints. Such constraints should not be altitudinal food-plant diversity variation either, as plant richness in fact decreased with our PC1 in the study area (Fig. 1 & Fig. S7). Thus, it is possible that certain non-food habitat properties co-varying with elevation, e.g., higher microhabitat variability (Fig. S15) or less-intense agriculture (with which our PC2 accounting only land cover could not fully capture), are important especially to butterflies and thus drive their local richness (Hill et al., 2021; Kleckova et al., 2014; Maes & Van Dyck, 2001). Conversely, for moths, food-plant diversity may be a more influential driver, as we saw congruently that plant richness increased with our PC2 (Fig. 1).

Regarding the potential diets of Lepidoptera across all constructed local networks, butterflies generally had narrower diets than the moths as larvae, but broader ones as adults. Note that these locally realised diets were determined by both the Lepidopterans’ biological diet breadths (feeding habits) and the local availabilities of food plants. We conducted a further comparison between butterflies’ and moths’ diets within our metawebs, i.e., accounting for only their biological diet breadths, and found consistent patterns (Fig. S14). Therefore, these patterns mainly mirrored biological diet-breadth difference among Lepidoptera clades. Butterfly larvae may experience comparatively stronger food-plant competition or predation, under which conditions narrower diet would be favoured (Dyer, 1995; Wiklund & Friberg, 2008). Meanwhile, butterfly adults tend to be more-generalist nectar feeders than moths likely because there are more plants flowering in day than in night (Borges et al., 2016). This has provided a broader spectrum of potential resources allowing the diurnal butterflies to evolve for adult food usage, whereas the fewer night-flowering plants and nocturnal moths have to more-tightly co-evolve. We also note that the strict diurnal vs. nocturnal active patterns between butterflies and moths, particularly at the adult stage, may have led to an underestimation of moths’ diets because observing their interactions with flowers are intrinsically more difficult in the dark. Nonetheless, since the compiled Lepidoptera-plant interaction information (see *Methods*) was long-term and extensive, it can be considered as near-complete (see also Pearse & Altermatt, 2015), and differences in adult diet breadth between butterflies and moths may not be an artefact only. Given that there is no systematic sampling bias expected along environmental gradients, the spatial patterns of Lepidoptera diet breadths (as well as of other metrics) should be realistic.

In general, local potential diet breadth of most Lepidoptera clades across their life stages tended to become broader toward low-elevation farmlands (low PC1 and high PC2), possibly because of more diverse plants in such habitats (Fig. 1) presenting more food plants to co-occur with the Lepidoptera. However, we cannot rule out that such local condition (more-perturbed farmlands) may favour the existence of generalist over specialist consumers (Büchi & Vuilleumier, 2014). An interesting exception was the adults of Geometrid moths, whose potential diets statistically broadened toward high-elevation forests (high PC1 and low PC2), countering the trend of local plant richness. It is possible that such conditions favour more-generalist Geometrids, driven by weaker plant resistance or other factors as reported in other plant-feeding insects (Moreira et al., 2018; Pellissier et al., 2012). However, given the reported Geometrids’ adult diets are narrow (Fig. S9), such trends were relatively trivial, and we remain unsure if there are mechanisms (as proposed) specific to Geometrid but not to other clades. The Lepidopterans’ local diet breadths were broader in observed communities than in randomly re-assembled ones across clades and life stages, except for adult Noctuid and Geometrid moths. The former pattern suggests generalist Lepidoptera to be relatively widespread (thus, per-grid re-assembling undersamples these generalists and leads to narrower diets), but this is less effective for adult moths who, despite variation in diet breadths, are generally narrow feeders. Relatively-generalist adult moths may be spatially restricted, or local plant assemblage tend to contain only a minor fraction of their food plants due to pollinator competition (thus per-grid re-assembling oversamples the generalists or their food plants and lead to broader diets). The latter condition could be shaped by pollinator competition among plants.

Network modularity emerges from groups of species interacting more within but less between the groups, which is typical in plant-herbivore and (large) plant-pollinator networks (Olesen et al., 2007; Thébault & Fontaine, 2010), including those of Lepidoptera larvae and adults (Astegiano et al., 2017). Evaluated with Lepidopterans’ local potential diets, butterflies’ larval networks were particularly modular whereas adult ones particularly non-modular over the moth clades. Part of such butterfly-moth difference should be associated with their diet-breadth difference as addressed above. As for butterflies, their diurnal adults embrace diverse flowering plants, and their nectar feeding is benefiting the plants with pollination functions. Under such conditions, co-evolution with plants would favour butterfly adults to feed on non-specialised, diverse nectars, thereby suppressing the formation of network modularity. Compare to the adults that can fly to access alternative nectar sources, their larvae are relatively immobile and thus needed to be capable to well-consume the individual host plant that they settle on. The larval co-evolutionary arm race with the host plants should consequently favour phylogenetically or physiologically conservative diets (Futuyma & Moreno, 1988), thereby making modularity prevail in larval networks (Fig. S10). Such inferences congruent with our null-model analyses, such that real-world Lepidoptera-plant interdependences (realistic butterfly diets) positively contributed to modularity in butterflies’ larval networks yet negatively in adult ones. As for moths, their adult networks were more modular than either type of null-model counterparts. This suggests a tighter co-evolution with food plants particularly in their adult phase, which is reasonable, given nocturnal flowering plants are just a handful.

As dietary modules within networks often emerge from narrow and specialized diets (Tim Tinker et al., 2012), it is expectable that the spatial-modularity patterns of Lepidoptera-plant networks should generally counter their spatial-generality patterns, which we indeed observed (Figs. 4 & 5; Figs. S9 & S10). On top of such associations, there are also mechanisms that can contribute to the formation of various modularity along environmental gradients. For example, the modularity of larval and adult butterfly networks were both negatively associated with farmland coverage (PC2), possibly due to anthropogenic influences. Agriculture modifies species composition of local communities to deviate from their phylogenetic structure or co-evolutionary legacy shaped before perturbation (Moora et al., 2014; Toyama et al., 2015), thereby mitigating the emergent network modularity. Consistently, as shown by null-model analyses, real-world species-environment reliance contributed negatively to butterfly networks’ modularity across life stages (more pronounced in adult ones) particularly toward high PC2. Meanwhile, the modularity of larval networks of Noctuid moths associated negatively with PC1. This pattern may reflect relatively more-effective local competitive exclusions toward high elevations (possible mechanisms see Montaño-Centellas et al., 2021), such that larvae sharing similar food plants (i.e., belonging to the same interaction module) tend not to co-occur.

In terms of potential dietary niche overlap in local networks, butterfly larvae had the lowest, while adults the highest, overlap among all clades. This echoes our above reasoning of diet specialisation of butterfly larvae vs. adults, as the same mechanisms lead also to differentiated diets among species. With larval networks, the environmental gradients were generally not influential to Lepidopterans’ niche overlap, while only butterfly larvae’s diets became less overlapped with PC1. Interestingly, if without other constraints, butterflies’ diets should passively become more overlapped given that their richness increases while plant richness decreases along PC1. Thus, such an inverse trend in larval networks may suggest a compositional shift toward dietary more-differentiated butterfly larvae at higher elevations (Pellissier et al., 2012), which should have suppressed their diet breath in addition to the effect of fewer food plants available. Contrastingly, environmental gradients were relatively influential to niche overlap in adult networks. The overlap tended to be higher toward low-elevation farmlands (low PC1 and high PC2) with butterflies and Noctuid moths, while toward high-elevation farmlands (high PC1 and PC2) with Geometrid moths, though the latter was relatively trivial. These patterns largely echo those of diet breadths and may be associated with the spatial pattern of plant richness, or the environmental conditions favouring dietary generalists over specialists. Given the particularly narrow diets of adult Geometrid moths, diet differentiation should be important for those locally coexisting. This is supported by that they adopted the lowest niche overlap among clades, and null models indicated that real-world constraints contributed negatively to their niche overlap.

By addressing the geo-climate and land-cover relevance of landscape-scale Lepidoptera-plant networks, we revealed across Lepidoptera clades and life stages how their (co-)evolved traits (e.g., biological diet breadth) and ecological relationships with others (e.g., resource competition, local food availability) collectively drive different interaction structures in response to environmental variations. Such biogeographical understandings provide direct and timely conservation implications given that the Lepidoptera in the study area are declining (Habel et al. 2019). For example, networks with high dietary overlap are more vulnerable to food-plant loss, whereas those with low modularity sensitive to cascading harmful perturbations (Pires et al., 2020; Stouffer & Bascompte, 2011). Identifying which local communities are of such, one can accordingly adjust management strategies for various aims, e.g., conserving specific pollinator species or the overall biodiversity. Moreover, the detected spatial patterns can inform how these communities tend to react to potential environmental changes caused by anthropogenic land use or climate change (Hill et al., 2021; Ho et al., 2022). We provided mechanistic explanations across evolutionary and ecological time scales for the detected network patterns, which could inform testable hypotheses. While here we compared Lepidoptera clades and life stages separately, in reality, they harbour in the same communities and influence each other. A taxonomically integrated revisit would be promising future work that can potentially provide further insights on the dynamical fate of Lepidoptera-plant communities via a network perspective.

## Acknowledgements

We thank all the many people who collected the occurrence data of Lepidoptera and plants upon which our analyses are based on, without their naturalist skills and efforts such analyses would simply not be possible. We specifically thank Robert Trusch und Arno Wörz (both Staatliches Museum für Naturkunde Stuttgart) for providing access to the Lepidoptera and plant data, respectively. We thank Rosi Siber for extracting needed GIS data. Funding is through the University of Zurich Research Priority Program in Global Change and Biodiversity (URPP GCB) as well as the Swiss National Science Foundation (Grant Nr. 310030_197410) to FA.

## Statement of author contributions

FA conceived the idea and developed it into a project together with HH. FA secured the funding. FA compiled the primary data and HH designed and performed the analyses. HH led the writing with consistent and significant input from FA.

## Data accessibility statement

The Lepidoptera and plant occurrence data can be obtained at *Database Arbeitsgruppe Schmetterlinge Baden-Württembergs, Bundesamt für Naturschutz* and *Floraweb*, respectively. The GIS data can be derived from corresponding databases (see Methods). The compiled metawebs can be accessed upon request to FA. The processed local network data, as well as the R codes that reproduce all analyses and figures in this study, will be provided at a public depository upon publication in a journal (or upon request to HH).

## References

Altermatt, F., & Pearse, I. S. (2011). Similarity and specialization of the larval versus adult diet of European butterflies and moths. The American Naturalist, 178(3), 372–382.

Astegiano, J., Altermatt, F., & Massol, F. (2017). Disentangling the co-structure of multilayer interaction networks: degree distribution and module composition in two-layer bipartite networks. Scientific Reports, 7(1), 1–11.

Axmacher, J. C., Brehm, G., Hemp, A., Tünte, H., Lyaruu, H. V., Müller-Hohenstein, K., & Fiedler, K. (2009). Determinants of diversity in afrotropical herbivorous insects (Lepidoptera: Geometridae): plant diversity, vegetation structure or abiotic factors? Journal of Biogeography, 36(2), 337–349.

Bascompte, J. (2009). Disentangling the web of life. Science, 325(5939), 416–419.

Beck, J., Liedtke, H. C., Widler, S., Altermatt, F., Loader, S. P., Hagmann, R., … & Fiedler, K. (2016). Patterns or mechanisms? Bergmann’s and Rapoport’s rule in moths along an elevational gradient. Community Ecology, 17(2), 137–148.

Boggs, C. L. (1987). Ecology of nectar and pollen feeding in Lepidoptera. Nutritional ecology of insects, mites, and spiders, 369–391.

Borges, R. M., Somanathan, H., & Kelber, A. (2016). Patterns and processes in nocturnal and crepuscular pollination services. The Quarterly Review of Biology, 91(4), 389–418.

Braga, M. P., Guimarães, P. R., Wheat, C. W., Nylin, S., & Janz, N. (2018). Unifying host-associated diversification processes using butterfly–plant networks. Nature communications, 9(1), 1–10.

Büchi, L., & Vuilleumier, S. (2014). Coexistence of specialist and generalist species is shaped by dispersal and environmental factors. The American Naturalist, 183(5), 612–624.

Carvajal Acosta, A. N., & Mooney, K. (2021). Effects of geographic variation in host plant resources for a specialist herbivore’s contemporary and future distribution. Ecosphere, 12(11), e03822.

Colwell, R. K., & Hurtt, G. C. (1994). Nonbiological gradients in species richness and a spurious Rapoport effect. The American Naturalist, 144(4), 570–595.

Core R Team (2013). R: A language and environment for statistical computing.

Després, L., David, J. P., & Gallet, C. (2007). The evolutionary ecology of insect resistance to plant chemicals. Trends in ecology & evolution, 22(6), 298–307.

Dormann, C. F., Fründ, J., Blüthgen, N., & Gruber, B. (2009). Indices, graphs and null models: analyzing bipartite ecological networks. The Open Ecology Journal, 2(1).

Dyer, L. A. (1995). Tasty generalists and nasty specialists? Antipredator mechanisms in tropical lepidopteran larvae. Ecology, 76(5), 1483–1496.

Ebert, G. (1991–2005). Die Schmetterlinge Baden-Württembergs. Vols. 1–10. Ulmer, Stuttgart.

Futuyma, D. J., & Moreno, G. (1988). The evolution of ecological specialization. Annual review of Ecology and Systematics, 207–233.

Gallou, A., Baillet, Y., Ficetola, G. F., & Després, L. (2017). Elevational gradient and human effects on butterfly species richness in the French Alps. Ecology and evolution, 7(11), 3672–3681.

Gibson, R. H., Nelson, I. L., Hopkins, G. W., Hamlett, B. J., & Memmott, J. (2006). Pollinator webs, plant communities and the conservation of rare plants: arable weeds as a case study. Journal of applied ecology, 43(2), 246–257.

Habel, J. C., Trusch, R., Schmitt, T., Ochse, M., & Ulrich, W. (2019). Long-term large-scale decline in relative abundances of butterfly and burnet moth species across south-western Germany. Scientific reports, 9(1), 1–9.

Hanspach, J., Schweiger, O., Kühn, I., Plattner, M., Pearman, P. B., Zimmermann, N. E., & Settele, J. (2014). Host plant availability potentially limits butterfly distributions under cold environmental conditions. Ecography, 37(3), 301–308.

Hill, G. M., Kawahara, A. Y., Daniels, J. C., Bateman, C. C., & Scheffers, B. R. (2021). Climate change effects on animal ecology: butterflies and moths as a case study. Biological Reviews, 96(5), 2113–2126.

Ho, H. C., Brodersen, J., Gossner, M. M., Graham, C. H., Kaeser, S., Chacko, M. R., … & Altermatt, F. (2022). Blue and green food webs respond differently to elevation and land use. Nature Communications (In Press).

Jones, J. P. (2011). Monitoring species abundance and distribution at the landscape scale. Journal of Applied Ecology, 48(1), 9–13.

Kawahara, A. Y., Plotkin, D., Espeland, M., Meusemann, K., Toussaint, E. F., Donath, A., … Breinholt, J. W. (2019). Phylogenomics reveals the evolutionary timing and pattern of butterflies and moths. Proceedings of the National Academy of Sciences, 116(45), 22657–22663.

Kleckova, I., Konvicka, M., & Klecka, J. (2014). Thermoregulation and microhabitat use in mountain butterflies of the genus Erebia: Importance of fine-scale habitat heterogeneity. Journal of Thermal Biology, 41, 50–58.

López-Carretero, A., Díaz-Castelazo, C., Boege, K., & Rico-Gray, V. (2014). Evaluating the spatiotemporal factors that structure network parameters of plant-herbivore interactions. PLoS One, 9(10), e110430.

Maes, D., & Van Dyck, H. (2001). Butterfly diversity loss in Flanders (north Belgium): Europe’s worst case scenario? Biological conservation, 99(3), 263–276.

Mantyka-Pringle, C. S., Visconti, P., Di Marco, M., Martin, T. G., Rondinini, C., & Rhodes, J. R. (2015). Climate change modifies risk of global biodiversity loss due to land-cover change. Biological Conservation, 187, 103–111.

Memmott, J. (1999). The structure of a plant-pollinator food web. Ecology letters, 2(5), 276–280.

Montaño-Centellas, F. A., Loiselle, B. A., & Tingley, M. W. (2021). Ecological drivers of avian community assembly along a tropical elevation gradient. Ecography, 44(4), 574–588.

Moora, M., Davison, J., Öpik, M., Metsis, M., Saks, Ü., Jairus, T., … & Zobel, M. (2014). Anthropogenic land use shapes the composition and phylogenetic structure of soil arbuscular mycorrhizal fungal communities. FEMS Microbiology Ecology, 90(3), 609–621.

Moreira, X., Petry, W. K., Mooney, K. A., Rasmann, S., & Abdala-Roberts, L. (2018). Elevational gradients in plant defences and insect herbivory: recent advances in the field and prospects for future research. Ecography, 41(9), 1485–1496.

Narango, D. L., Tallamy, D. W., & Shropshire, K. J. (2020). Few keystone plant genera support the majority of Lepidoptera species. Nature communications, 11(1), 1–8.

Olesen, J. M., Bascompte, J., Dupont, Y. L., & Jordano, P. (2007). The modularity of pollination networks. Proceedings of the National Academy of Sciences, 104(50), 19891–19896.

Pearse, I. S., & Altermatt, F. (2013a). Predicting novel trophic interactions in a non-native world. Ecology letters, 16(8), 1088–1094.

Pearse, I. S., & Altermatt, F. (2013b). Extinction cascades partially estimate herbivore losses in a complete Lepidoptera–plant food web. Ecology, 94(8), 1785–1794.

Pearse, I. S., & Altermatt, F. (2015). Out-of-sample predictions from plant–insect food webs: robustness to missing and erroneous trophic interaction records. Ecological Applications, 25(7), 1953–1961.

Pellissier, L., Albouy, C., Bascompte, J., Farwig, N., Graham, C., Loreau, M., … Gravel, D. (2018). Comparing species interaction networks along environmental gradients. Biological Reviews, 93(2), 785–800.

Pellissier, L., Fiedler, K., Ndribe, C., Dubuis, A., Pradervand, J. N., Guisan, A., & Rasmann, S. (2012). Shifts in species richness, herbivore specialization, and plant resistance along elevation gradients. Ecology and Evolution, 2(8), 1818–1825.

Pires, M. M., O’Donnell, J. L., Burkle, L. A., Díaz-Castelazo, C., Hembry, D. H., Yeakel, J. D., … Guimarães Jr, P. R. (2020). The indirect paths to cascading effects of extinctions in mutualistic networks.

Saravia, L. A., Marina, T. I., Kristensen, N. P., De Troch, M., & Momo, F. R. (2022). Ecological network assembly: how the regional metaweb influences local food webs. Journal of Animal Ecology, 91(3), 630–642.

Scoble, M. J. (1995). The Lepidoptera. Form, function and diversity. Oxford University Press.

Stouffer, D. B., & Bascompte, J. (2011). Compartmentalization increases food-web persistence. Proceedings of the National Academy of Sciences, 108(9), 3648–3652.

thébault, E., & Fontaine, C. (2010). Stability of ecological communities and the architecture of mutualistic and trophic networks. Science, 329(5993), 853–856.

tim Tinker, M., Guimaraes Jr, P. R., Novak, M., Marquitti, F. M. D., Bodkin, J. L., Staedler, M., … & Estes, J. A. (2012). Structure and mechanism of diet specialisation: testing models of individual variation in resource use with sea otters. Ecology letters, 15(5), 475–483.

thompson, R. M., Brose, U., Dunne, J. A., Hall Jr, R. O., Hladyz, S., Kitching, R. L., … Tylianakis, J. M. (2012). Food webs: reconciling the structure and function of biodiversity. Trends in ecology & evolution, 27(12), 689–697.

toyama, H., Kajisa, T., Tagane, S., Mase, K., Chhang, P., Samreth, V., … & Yahara, T. (2015). Effects of logging and recruitment on community phylogenetic structure in 32 permanent forest plots of Kampong Thom, Cambodia. Philosophical Transactions of the Royal Society B: Biological Sciences, 370(1662), 20140008.

tylianakis, J. M., Laliberté, E., Nielsen, A., & Bascompte, J. (2010). Conservation of species interaction networks. Biological conservation, 143(10), 2270–2279.

tylianakis, J. M., & Morris, R. J. (2017). Ecological networks across environmental gradients. Annual Review of Ecology, Evolution, and Systematics, 48, 25–48.

Weiblen, G. D., Webb, C. O., Novotny, V., Basset, Y., & Miller, S. E. (2006). Phylogenetic dispersion of host use in a tropical insect herbivore community. Ecology, 87(sp7), S62–S75.

Wiklund, C., & Friberg, M. (2008). Enemy-free space and habitat-specific host specialization in a butterfly. Oecologia, 157(2), 287–294.

