## Supplementary Information for "The structure of Lepidoptera-plant interaction networks across clades, life stages, and environmental gradients"

### Section S1. Additional details of network quantification and analyses

Besides the three focal metrics (i.e., mean generality, modularity, and niche overlap) mentioned in the main text, we also quantified the connectance and nestedness of the constructed Lepidoptera-plant networks. Connectance reflects how connected the local network is by considering the total number of realised links standardised by the number of species. Nestedness indicates the prevalence of a nested structure within a network, such that the therein (interaction) specialists tend to interact with a subset of partners of the generalists (Thébault & Fontaine, 2010).

All network metric quantification and analyses were done in R language. For quantifying mean generality, we summed up the number of food plant per Lepidoptera species with a given constructed local network (of the focal Lepidoptera clade and stage; the same for all metric calculation below), then averaged across all Lepidoptera species. For connectance, with a given local network, we calculated its total number of links divided by its total potential links, i.e., the product of number of plant species and Lepidoptera species in this given network. For nestedness, we used the nestednodf function from the vegan package v. 2.6-2 (Oksanen et al., 2022). For modularity, we used the metaComputeModules function (with "Beckett" algorithm) from the bipartite package v. 2.17 (Dormann et al., 2009). For dietary niche overlap of the consumers, we adopted the inbuilt Horn's index calculation of the networklevel function from the bipartite package. We generated null-model counterpart networks with matrix operations also in R (see explanation of the null models in the main text), then quantified them as for the constructed local networks (observations) as above described. To convert the observations to Z scores, we took the difference between observed value and the mean of the null-model values (former minus latter) then divided it by the standard deviation of the latter.

We carried out principal component analyses of environmental variables using R-base prcomp function, then visualised the result with the ggbiplot function from the ggbiplot package (Vu, 2011). We conducted general linear model analyses and visualized corresponding 2D and 3D regressions using the inbuilt lm function of the ggplot2 package v. 3.3.6 (Wickham, 2016), and a combination of R-base lm function with the plot_ly function (plotly package v. 4.10.0; Sievert, 2020), respectively. The ggpubr (v. 0.4.0; Kassabara, 2020), ggpmisc (v. 0.4.7; Aphalo, 2020), and gridExtra (v. 2.3; Auguie, 2017) packages were applied with ggplot2 for generating needed components of the plots. We illustrated the location and composition of local networks on a Baden-Württemberg area frame using the geom_scatterpie function from the scatterpie package v. 0.1.7 (Yu, 2021) and function png from the png package v 0.1-7 (Urbanek, 2013). We generated the correlation matrix between Lepidoptera clades and plants using the ggpairs function of the GGally package v. 2.1.2 (Schloerke et al., 2021).

All relevant R codes are available at the provided repository.

**Section S2. Additional figures and tables**

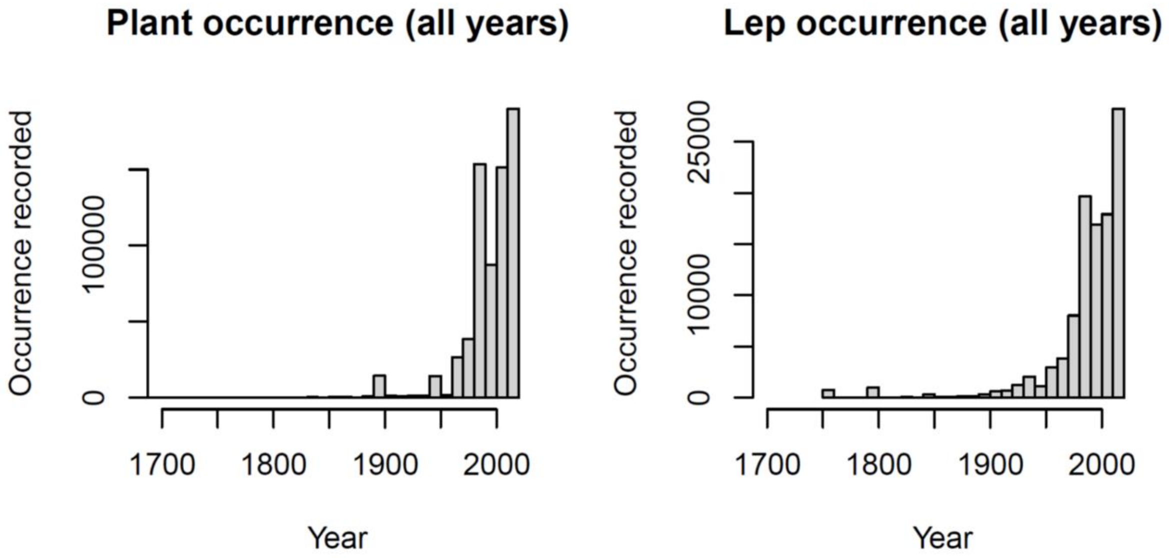


Figure S1. The yearly number of records (collapsing species and sites) of plant (left) and Lepidoptera (right) occurrence throughout our compiled data. In this study, we use occurrence information from the three decades between 1985 and 2014, which are represented by (roughly) the rightmost four bars on both plots.

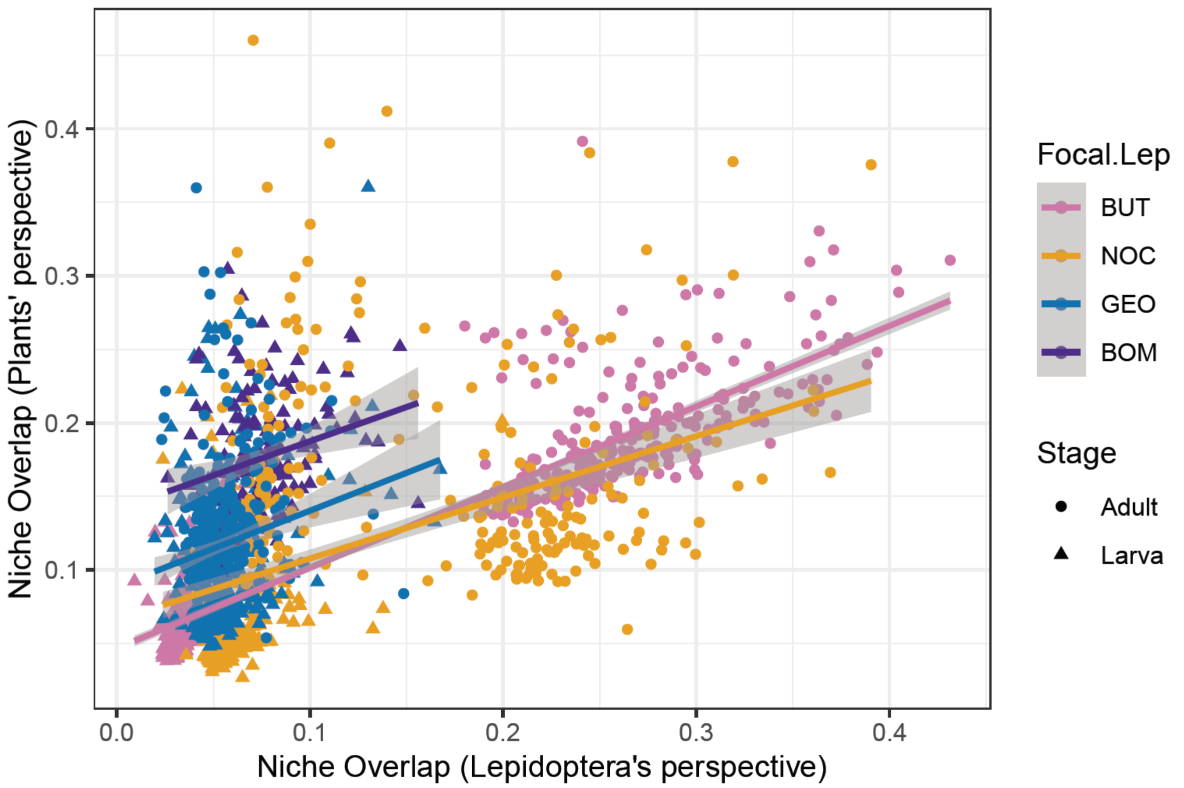


Figure S2. Positive associations between the niche overlap indices evaluated from the Lepidoptera's perspective vs. the plants' perspective across all the local networks we studied. The shades indicate respective 95% CI.

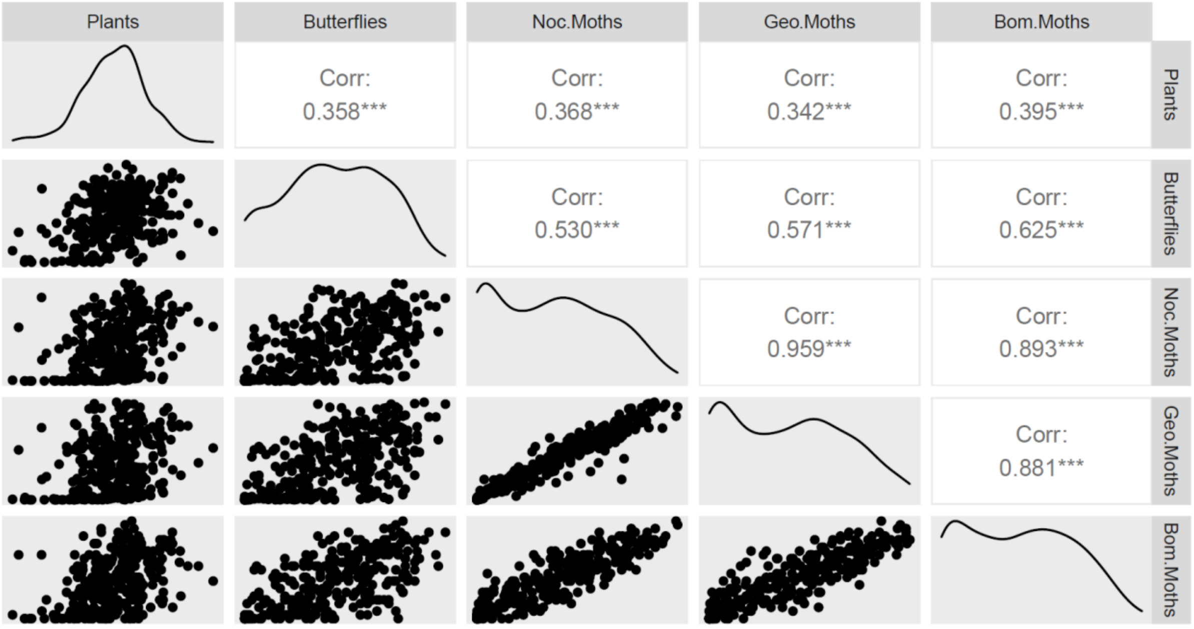


Figure S3. The correlation matrix among local (per grid, N = 310) richness of plants and the four focal clades of Lepidoptera, namely butterflies, Noctuid (Noc.) moths, Geometrid (Geo.) moths, and Bombycoidea (Bom.) moths. The lower triangle of the matrix shows the scatter plots of each combination of richness comparison, the upper triangle the corresponding Pearson correlation coefficients (all positive and with $p < 0.001$), and diagonal the density curves.

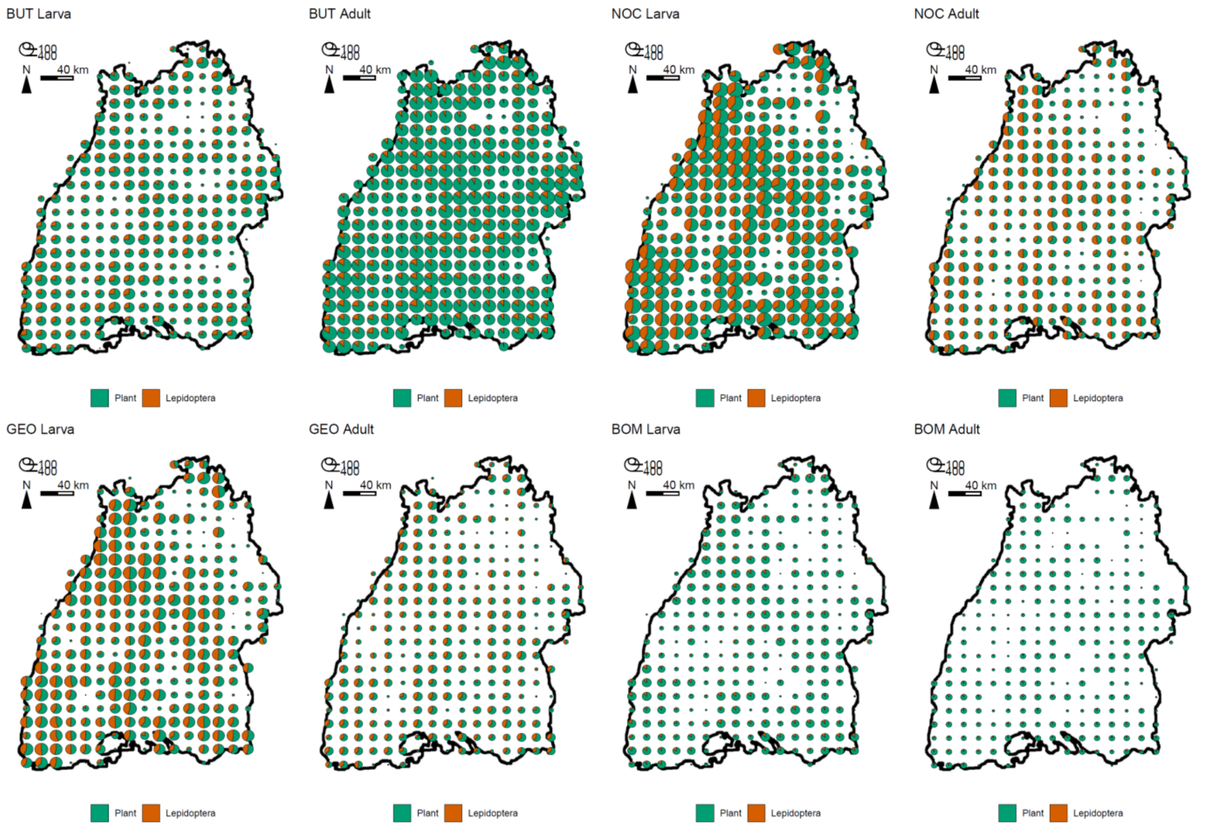


Figure S4. The size (number of nodes) and composition (Plants vs. Lepidoptera) of Lepidoptera-plant network in each grid across Baden-Württemberg. The four focal clades of Lepidoptera, namely butterflies (BUT), Noctuid moths (NOC), Geometrid moths (GEO), and Bombycoid moths (BOM), and their two distinct life stages, are presented in separate subfigures.

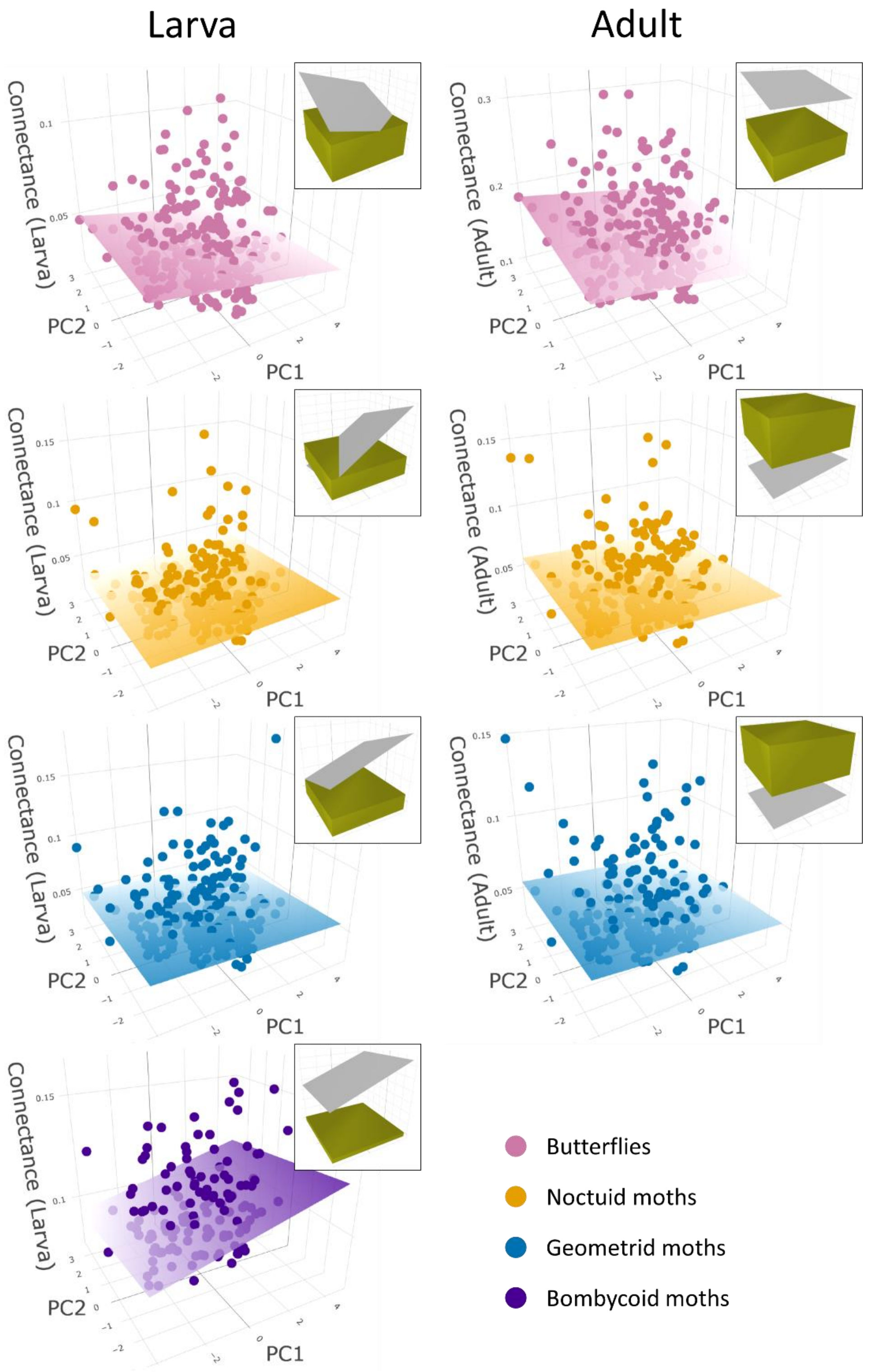


Figure S5. Connectance of the local networks across environmental gradients as 3D scatterplots, sharing the same environmental PC axes (as in main text Fig. 2) but respective metric reading axis. The planes in the 3D scatterplots are regression planes of observed values, where their colour fade toward the low-value end. Corresponding stats with separated regressions against each PC are provided in Fig. S12. Corresponding null-model analysis against randomised counterparts is given at each subplot's top-right (details see main text *Methods* and Fig. 3). Note that since the total number of nodes and links, and thus connectance, were fixed to empirical values in the "shuffled" null model, only the "re-assembled" planes appear in the null-model analyses.

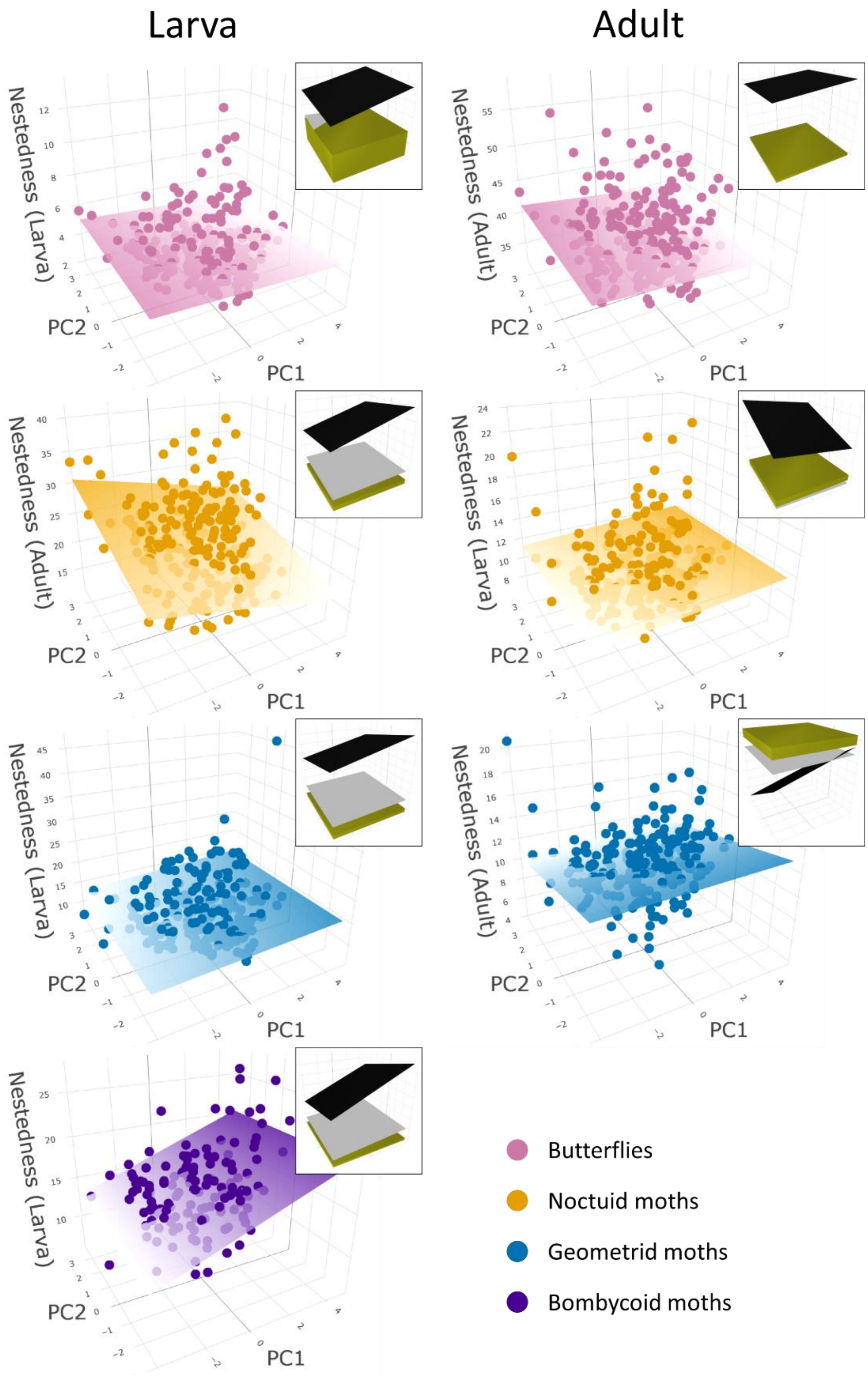


Figure S6. Nestedness of the local networks across environmental gradients as 3D scatterplots, sharing the same environmental PC axes (as in main text Fig. 2) but respective metric reading axis. The planes in the 3D scatterplots are regression planes of observed values, where their colour fade toward the low-value end. Corresponding stats with separated regressions against each PC are provided in Fig. S13. Corresponding null-model analysis against randomised counterparts is given at each subplot's top-right (details see main text *Methods* and Fig. 3).

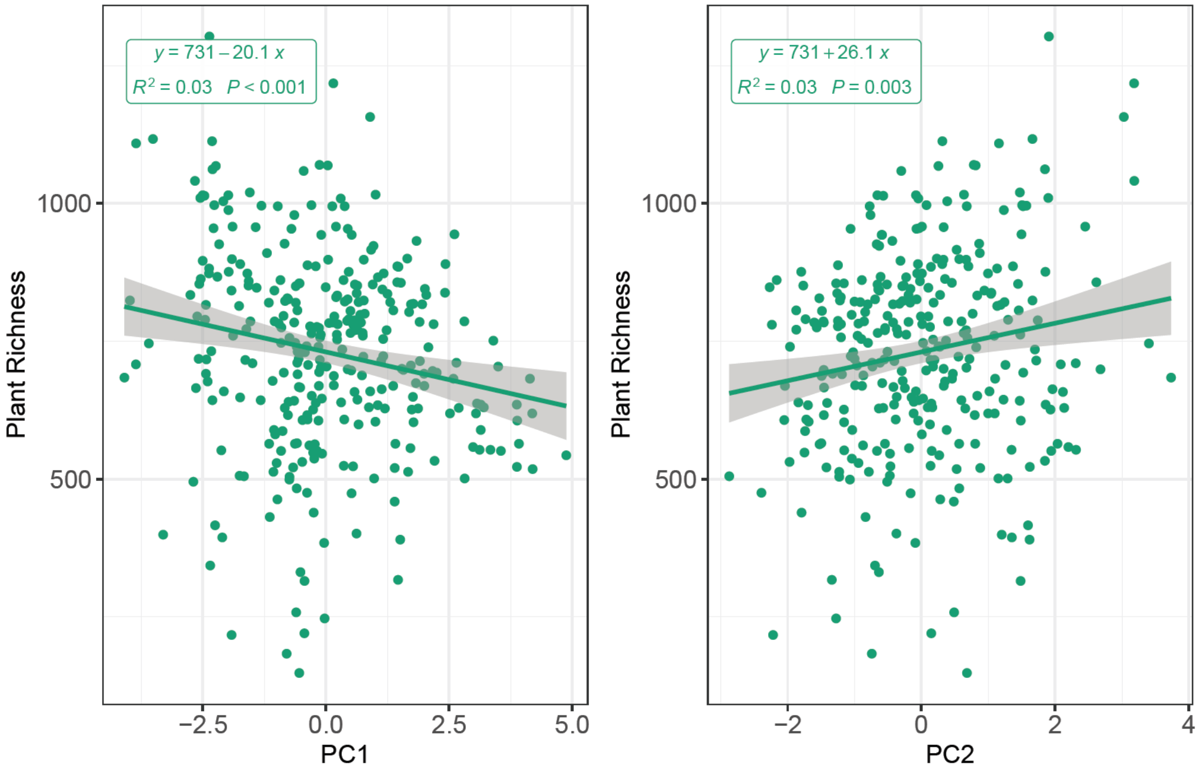


Figure S7. Plant richness against respective PC1 (left panel) and PC2 (middle panel) with our compiled occurrence data. The regression lines with SE (shades) are presented alongside corresponding regression equations and P values, where a P < 0.05 indicates a significant non-zero slope (i.e., the PC is influential).

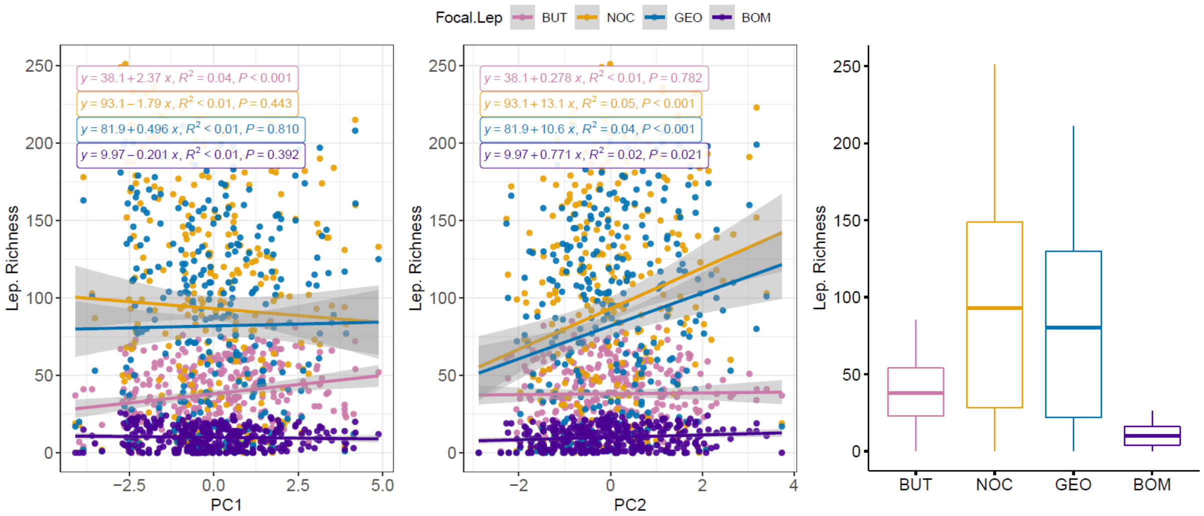


| | Df | Sum Sq | Mean Sq | F value | Pr(>F) |
|---|---|---|---|---|---|
| *Focal Lep.* | 3 | 1391493 | 463831.062 | 206.115 | < 0.001 |
| *Residuals* | 1236 | 2781438 | 2250.354 | NA | NA |

| | Df | Sum Sq | Mean Sq | F value | Pr(>F) |
|---|---|---|---|---|---|
| *Focal Lep.* | 3 | 1391493.187 | 463831.062 | 206.044 | < 0.001 |
| *PC1* | 1 | 170.135 | 170.135 | 0.076 | 0.783 |
| *Focal Lep. × PC1* | 3 | 7873.730 | 2624.577 | 1.166 | 0.322 |
| *Residuals* | 1232 | 2773393.721 | 2251.131 | NA | NA |

| | Df | Sum Sq | Mean Sq | F value | Pr(>F) |
|---|---|---|---|---|---|
| *Focal Lep.* | 3 | 1391493.19 | 463831.062 | 215.034 | < 0.001 |
| *PC2* | 1 | 66682.09 | 66682.086 | 30.914 | < 0.001 |
| *Focal Lep. × PC2* | 3 | 57310.35 | 19103.451 | 8.856 | < 0.001 |
| *Residuals* | 1232 | 2657445.15 | 2157.017 | NA | NA |

| | diff | lwr | upr | p adj |
|---|---|---|---|---|
| *NOC−BUT* | 55.026 | 45.224 | 64.828 | < 0.001 |
| *GEO−BUT* | 43.813 | 34.011 | 53.615 | < 0.001 |
| *BOM−BUT* | −28.129 | −37.931 | −18.327 | < 0.001 |
| *GEO−NOC* | −11.213 | −21.015 | −1.411 | 0.017 |
| *BOM−NOC* | −83.155 | −92.957 | −73.353 | < 0.001 |
| *BOM−GEO* | −71.942 | −81.744 | −62.140 | < 0.001 |

Figure S8. Lepidoptera richness of the four focal clades (butterflies: BUT; Noctuid moths: NOC; Geometrid moths: GEO; Bombycoid moths: BOM), regressed using GLM against respective PC1 (left panel) and PC2 (middle panel), and as a regional overall among-clade comparison (right panel; effectively an ANOVA). Corresponding stats are given in stats tables below the plots. In the former two panels, the regression lines with SE (shades) are presented alongside corresponding regression equations and P values grouped by colours, where a $P < 0.05$ indicates a significant non-zero slope (i.e., the PC is influential). A $P < 0.05$ in the metric * PC term indicates different slopes among Lepidoptera clades. The stats table at the right presents whether there is significant among-clade difference when looking at the whole region, and if yes, which exact pairs have such differences. Comparing to the 3D scatterplots in the main text, the left and middle plots can be seen as collapsing the PC2 and PC1 axis, respectively, while the right plot as collapsing both PC axes.

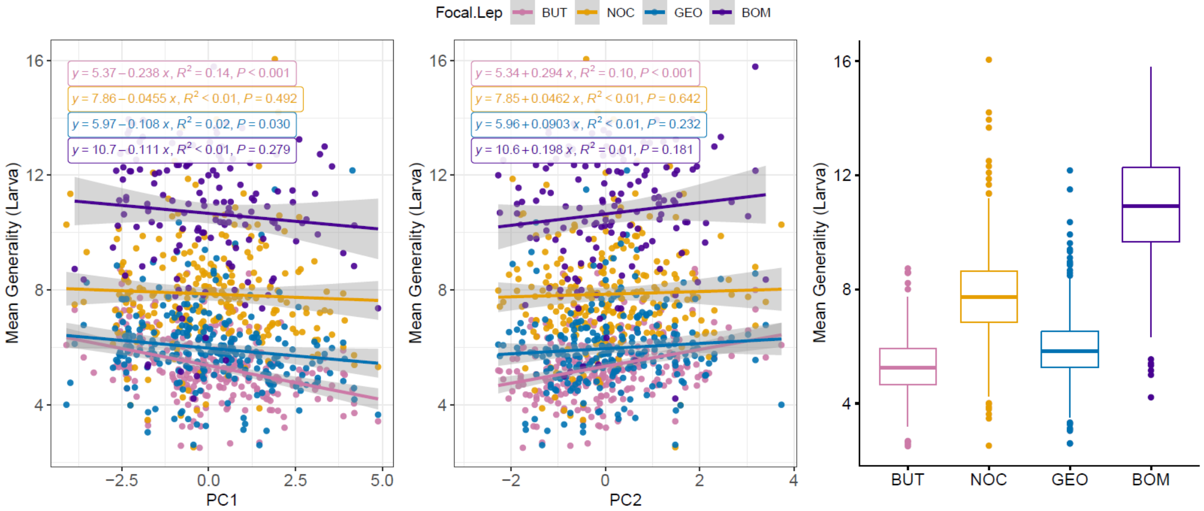


| | Df | Sum Sq | Mean Sq | F value | Pr(>F) |
|---|---|---|---|---|---|
| *Focal Lep.* | 3 | 3173.766 | 1057.922 | 432.149 | < 0.001 |
| *PC1* | 1 | 45.472 | 45.472 | 18.575 | < 0.001 |
| *Focal Lep. × PC1* | 3 | 15.215 | 5.072 | 2.072 | 0.102 |
| *Residuals* | 913 | 2235.066 | 2.448 | NA | NA |

| | Df | Sum Sq | Mean Sq | F value | Pr(>F) |
|---|---|---|---|---|---|
| *Focal Lep.* | 3 | 3173.766 | 1057.922 | 428.895 | < 0.001 |
| *PC2* | 1 | 30.754 | 30.754 | 12.468 | < 0.001 |
| *Focal Lep. × PC2* | 3 | 12.975 | 4.325 | 1.753 | 0.155 |
| *Residuals* | 913 | 2252.025 | 2.467 | NA | NA |

| | Df | Sum Sq | Mean Sq | F value | Pr(>F) |
|---|---|---|---|---|---|
| *Focal Lep.* | 3 | 3173.766 | 1057.922 | 422.569 | < 0.001 |
| *Residuals* | 917 | 2295.753 | 2.504 | NA | NA |

| | diff | lwr | upr | p adj |
|---|---|---|---|---|
| *NOC−BUT* | 2.515 | 2.159 | 2.871 | < 0.001 |
| *GEO−BUT* | 0.624 | 0.268 | 0.980 | < 0.001 |
| *BOM−BUT* | 5.335 | 4.917 | 5.753 | < 0.001 |
| *GEO−NOC* | −1.891 | −2.254 | −1.528 | < 0.001 |
| *BOM−NOC* | 2.820 | 2.396 | 3.244 | < 0.001 |
| *BOM−GEO* | 4.711 | 4.288 | 5.135 | < 0.001 |

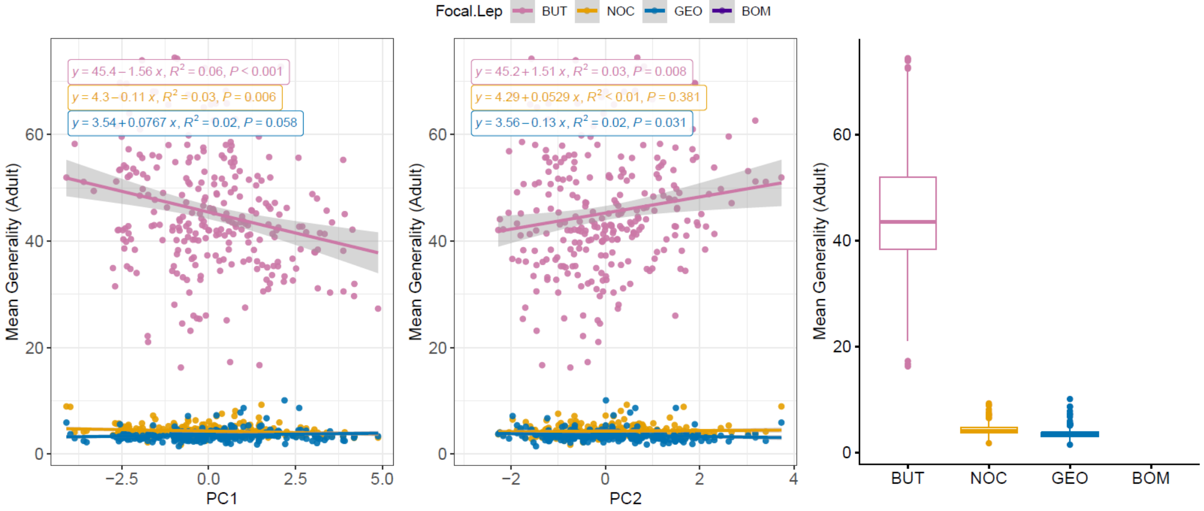


| | Df | Sum Sq | Mean Sq | F value | Pr(>F) |
|---|---|---|---|---|---|
| *Focal Lep.* | 2 | 294993.486 | 147496.743 | 3466.178 | < 0.001 |
| *PC1* | 1 | 701.848 | 701.848 | 16.493 | < 0.001 |
| *Focal Lep. × PC1* | 2 | 1237.894 | 618.947 | 14.545 | < 0.001 |
| *Residuals* | 740 | 31489.318 | 42.553 | NA | NA |

| | Df | Sum Sq | Mean Sq | F value | Pr(>F) |
|---|---|---|---|---|---|
| *Focal Lep.* | 2 | 294993.486 | 147496.743 | 3351.173 | < 0.001 |
| *PC2* | 1 | 282.449 | 282.449 | 6.417 | 0.012 |
| *Focal Lep. × PC2* | 2 | 576.648 | 288.324 | 6.551 | 0.002 |
| *Residuals* | 740 | 32569.963 | 44.013 | NA | NA |

| | Df | Sum Sq | Mean Sq | F value | Pr(>F) |
|---|---|---|---|---|---|
| *Focal Lep.* | 2 | 294993.49 | 147496.743 | 3278.288 | < 0.001 |
| *Residuals* | 743 | 33429.06 | 44.992 | NA | NA |

| | diff | lwr | upr | p adj |
|---|---|---|---|---|
| *NOC−BUT* | −40.966 | −42.363 | −39.568 | < 0.001 |
| *GEO−BUT* | −41.724 | −43.126 | −40.321 | < 0.001 |
| *GEO−NOC* | −0.758 | −2.204 | 0.687 | 0.435 |

Figure S9. GLM analyses on mean generality of the four focal clades, along PC1 (left panel) and PC2 (middle panel), or as a regional overall among-clade comparison (right panel), with life stages separated (upper: larva; lower: adult). See Fig. S8 for annotation explanations and detailed interpretation.

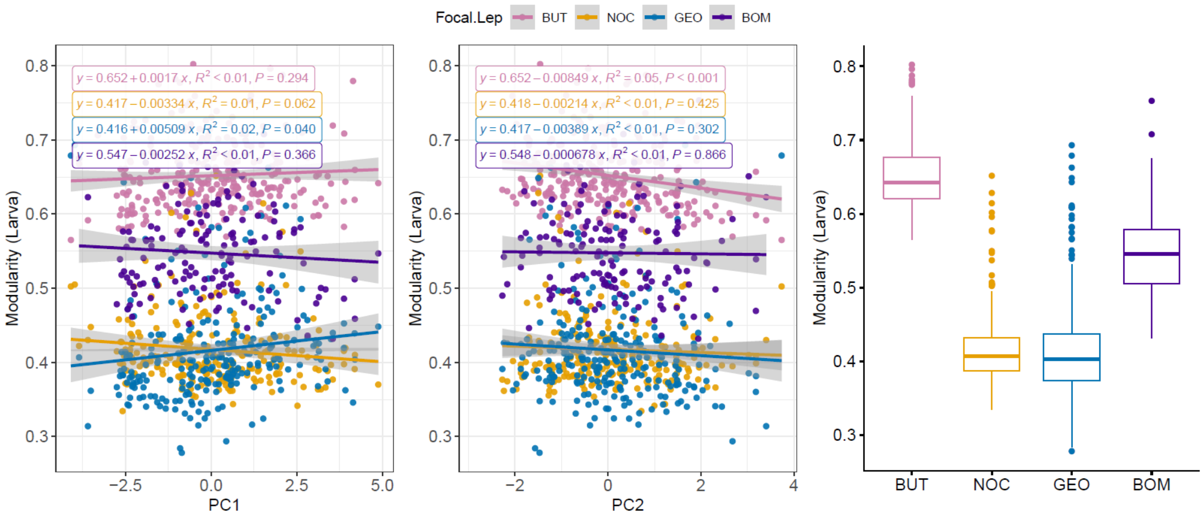


| | Df | Sum Sq | Mean Sq | F value | Pr(>F) |
|---|---|---|---|---|---|
| *Focal Lep.* | 3 | 10.032 | 3.344 | 1084.301 | < 0.001 |
| *PC1* | 1 | 0.001 | 0.001 | 0.335 | 0.563 |
| *Focal Lep. × PC1* | 3 | 0.033 | 0.011 | 3.526 | 0.015 |
| *Residuals* | 913 | 2.816 | 0.003 | NA | NA |

| | Df | Sum Sq | Mean Sq | F value | Pr(>F) |
|---|---|---|---|---|---|
| *Focal Lep.* | 3 | 10.032 | 3.344 | 1084.283 | < 0.001 |
| *PC2* | 1 | 0.023 | 0.023 | 7.379 | 0.007 |
| *Focal Lep. × PC2* | 3 | 0.011 | 0.004 | 1.173 | 0.319 |
| *Residuals* | 913 | 2.816 | 0.003 | NA | NA |

| | Df | Sum Sq | Mean Sq | F value | Pr(>F) |
|---|---|---|---|---|---|
| *Focal Lep.* | 3 | 10.032 | 3.344 | 1076.187 | < 0.001 |
| *Residuals* | 917 | 2.849 | 0.003 | NA | NA |

| | diff | lwr | upr | p adj |
|---|---|---|---|---|
| *NOC−BUT* | −0.234 | −0.247 | −0.222 | < 0.001 |
| *GEO−BUT* | −0.235 | −0.248 | −0.223 | < 0.001 |
| *BOM−BUT* | −0.104 | −0.119 | −0.090 | < 0.001 |
| *GEO−NOC* | −0.001 | −0.014 | 0.012 | 0.997 |
| *BOM−NOC* | 0.130 | 0.115 | 0.145 | < 0.001 |
| *BOM−GEO* | 0.131 | 0.116 | 0.146 | < 0.001 |

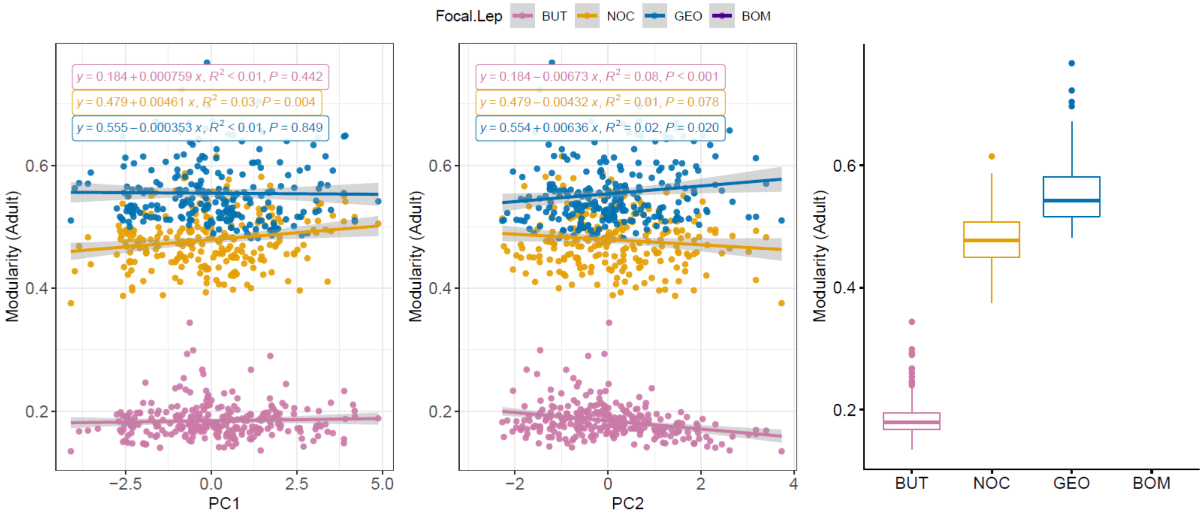


| | Df | Sum Sq | Mean Sq | F value | Pr(>F) |
|---|---|---|---|---|---|
| *Focal Lep.* | 2 | 19.739 | 9.869 | 5913.520 | < 0.001 |
| *PC1* | 1 | 0.006 | 0.006 | 3.685 | 0.055 |
| *Focal Lep. × PC1* | 2 | 0.010 | 0.005 | 2.992 | 0.051 |
| *Residuals* | 740 | 1.235 | 0.002 | NA | NA |

| | Df | Sum Sq | Mean Sq | F value | Pr(>F) |
|---|---|---|---|---|---|
| *Focal Lep.* | 2 | 19.739 | 9.869 | 6011.050 | < 0.001 |
| *PC2* | 1 | 0.003 | 0.003 | 1.936 | 0.165 |
| *Focal Lep. × PC2* | 2 | 0.033 | 0.016 | 10.048 | < 0.001 |
| *Residuals* | 740 | 1.215 | 0.002 | NA | NA |

| | Df | Sum Sq | Mean Sq | F value | Pr(>F) |
|---|---|---|---|---|---|
| *Focal Lep.* | 2 | 19.739 | 9.869 | 5860.919 | < 0.001 |
| *Residuals* | 743 | 1.251 | 0.002 | NA | NA |

| | diff | lwr | upr | p adj |
|---|---|---|---|---|
| *NOC−BUT* | 0.295 | 0.286 | 0.303 | < 0.001 |
| *GEO−BUT* | 0.371 | 0.362 | 0.379 | < 0.001 |
| *GEO−NOC* | 0.076 | 0.067 | 0.085 | < 0.001 |

Figure S10. GLM analyses on network modularity of the four focal clades, along PC1 (left panel) and PC2 (middle panel), or as a regional overall among-clade comparison (right panel), with life stages separated (upper: larva; lower: adult). See Fig. S8 for annotation explanations and detailed interpretation.

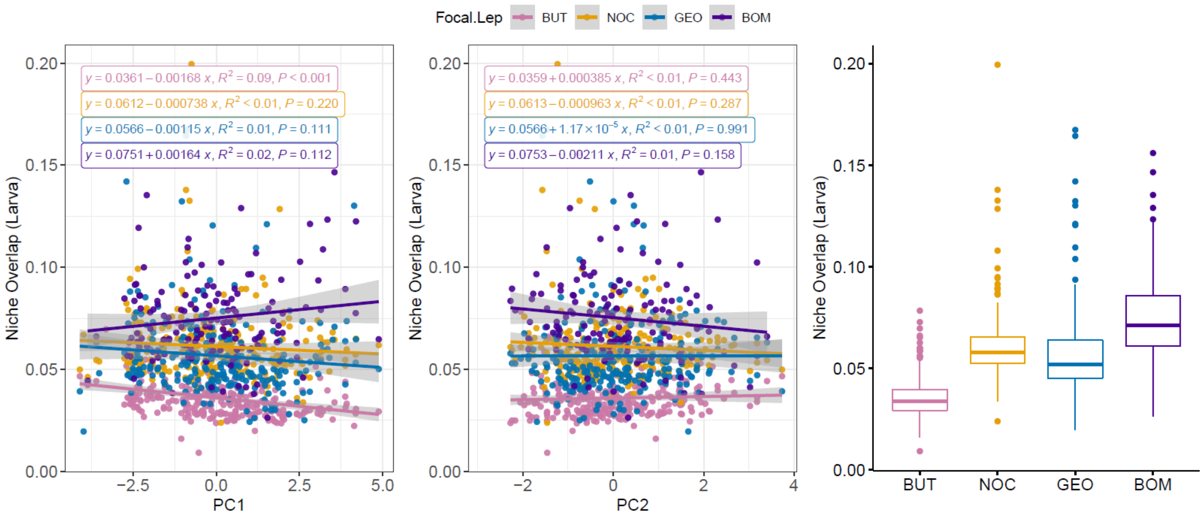


| | Df | Sum Sq | Mean Sq | F value | Pr(>F) |
|---|---|---|---|---|---|
| *Focal Lep.* | 3 | 0.168 | 0.056 | 197.573 | < 0.001 |
| *PC1* | 1 | 0.002 | 0.002 | 5.501 | 0.019 |
| *Focal Lep. × PC1* | 3 | 0.003 | 0.001 | 3.848 | 0.009 |
| *Residuals* | 913 | 0.258 | 0.000 | NA | NA |

| | Df | Sum Sq | Mean Sq | F value | Pr(>F) |
|---|---|---|---|---|---|
| *Focal Lep.* | 3 | 0.168 | 0.056 | 194.907 | < 0.001 |
| *PC2* | 1 | 0.000 | 0.000 | 1.043 | 0.308 |
| *Focal Lep. × PC2* | 3 | 0.001 | 0.000 | 1.150 | 0.328 |
| *Residuals* | 913 | 0.262 | 0.000 | NA | NA |

| | Df | Sum Sq | Mean Sq | F value | Pr(>F) |
|---|---|---|---|---|---|
| *Focal Lep.* | 3 | 0.168 | 0.056 | 194.802 | < 0.001 |
| *Residuals* | 917 | 0.263 | 0.000 | NA | NA |

| | diff | lwr | upr | p adj |
|---|---|---|---|---|
| *NOC–BUT* | 0.025 | 0.021 | 0.029 | < 0.001 |
| *GEO–BUT* | 0.021 | 0.017 | 0.024 | < 0.001 |
| *BOM–BUT* | 0.039 | 0.035 | 0.044 | < 0.001 |
| *GEO–NOC* | −0.005 | −0.008 | −0.001 | 0.013 |
| *BOM–NOC* | 0.014 | 0.009 | 0.018 | < 0.001 |
| *BOM–GEO* | 0.018 | 0.014 | 0.023 | < 0.001 |

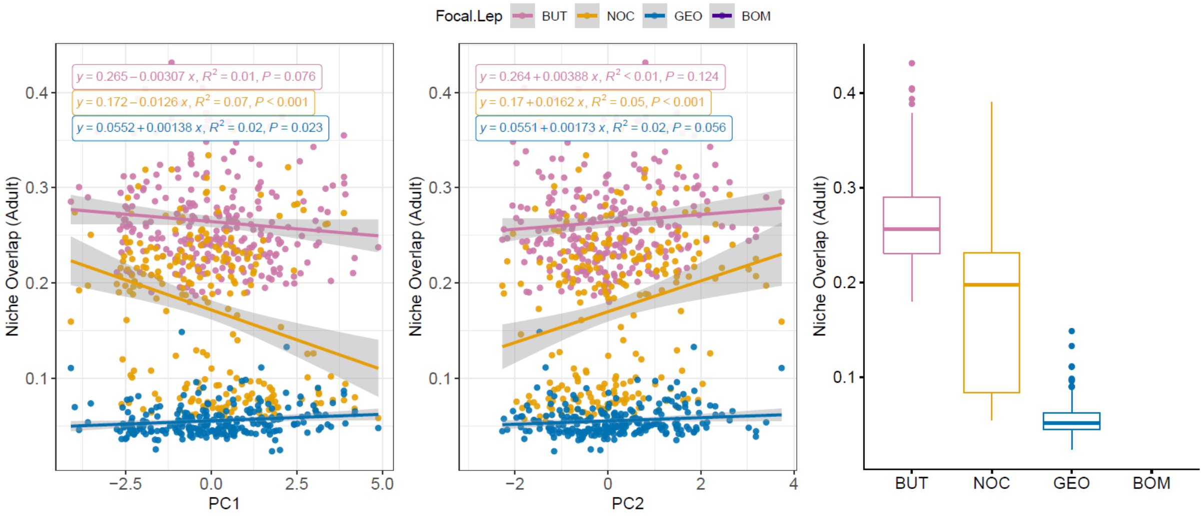


| | Df | Sum Sq | Mean Sq | F value | Pr(>F) |
|---|---|---|---|---|---|
| *Focal Lep.* | 2 | 5.508 | 2.754 | 939.901 | < 0.001 |
| *PC1* | 1 | 0.050 | 0.050 | 17.174 | < 0.001 |
| *Focal Lep. × PC1* | 2 | 0.075 | 0.037 | 12.729 | < 0.001 |
| *Residuals* | 740 | 2.168 | 0.003 | NA | NA |

| | Df | Sum Sq | Mean Sq | F value | Pr(>F) |
|---|---|---|---|---|---|
| *Focal Lep.* | 2 | 5.508 | 2.754 | 925.566 | < 0.001 |
| *PC2* | 1 | 0.051 | 0.051 | 17.198 | < 0.001 |
| *Focal Lep. × PC2* | 2 | 0.040 | 0.020 | 6.749 | 0.001 |
| *Residuals* | 740 | 2.202 | 0.003 | NA | NA |

| | Df | Sum Sq | Mean Sq | F value | Pr(>F) |
|---|---|---|---|---|---|
| *Focal Lep.* | 2 | 5.508 | 2.754 | 892.306 | < 0.001 |
| *Residuals* | 743 | 2.293 | 0.003 | NA | NA |

| | diff | lwr | upr | p adj |
|---|---|---|---|---|
| *NOC–BUT* | −0.092 | −0.104 | −0.081 | < 0.001 |
| *GEO–BUT* | −0.209 | −0.220 | −0.197 | < 0.001 |
| *GEO–NOC* | −0.117 | −0.129 | −0.105 | < 0.001 |

Figure S11. GLM analyses on dietary niche overlap of the four focal clades, along PC1 (left panel) and PC2 (middle panel), or as a regional overall among-clade comparison (right panel), with life stages separated (upper: larva; lower: adult). See Fig. S8 for annotation explanations and detailed interpretation.

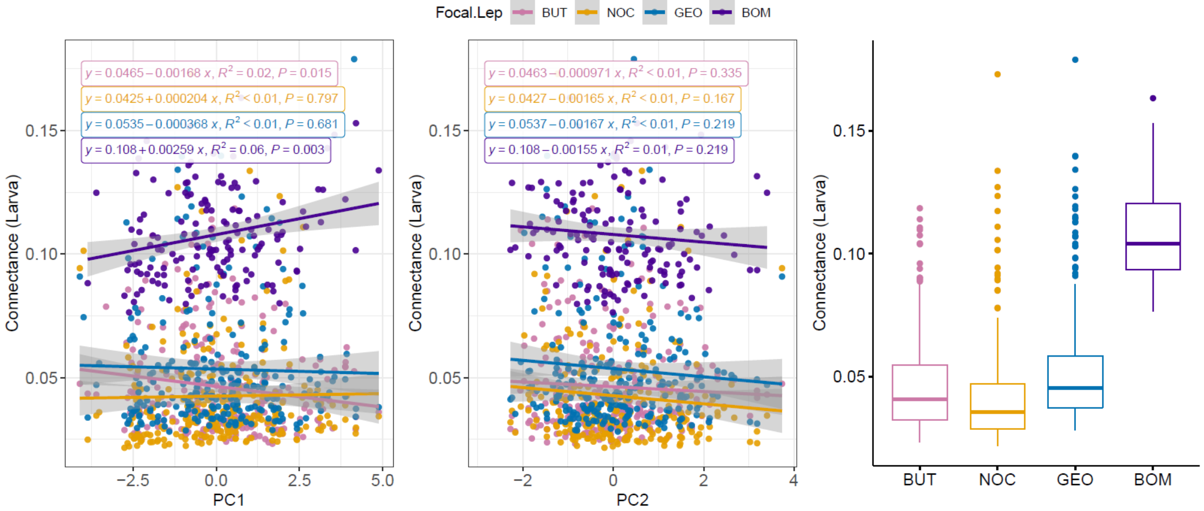


| | Df | Sum Sq | Mean Sq | F value | Pr(>F) |
|---|---|---|---|---|---|
| *Focal Lep.* | 3 | 0.462 | 0.154 | 333.274 | < 0.001 |
| *Residuals* | 917 | 0.424 | 0.000 | NA | NA |

| | Df | Sum Sq | Mean Sq | F value | Pr(>F) |
|---|---|---|---|---|---|
| *Focal Lep.* | 3 | 0.462 | 0.154 | 335.974 | < 0.001 |
| *PC1* | 1 | 0.000 | 0.000 | 0.091 | 0.763 |
| *Focal Lep. × PC1* | 3 | 0.005 | 0.002 | 3.779 | 0.01 |
| *Residuals* | 913 | 0.419 | 0.000 | NA | NA |

| | Df | Sum Sq | Mean Sq | F value | Pr(>F) |
|---|---|---|---|---|---|
| *Focal Lep.* | 3 | 0.462 | 0.154 | 333.946 | < 0.001 |
| *PC2* | 1 | 0.003 | 0.003 | 5.597 | 0.018 |
| *Focal Lep. × PC2* | 3 | 0.000 | 0.000 | 0.084 | 0.969 |
| *Residuals* | 913 | 0.421 | 0.000 | NA | NA |

| | diff | lwr | upr | p adj |
|---|---|---|---|---|
| *NOC−BUT* | −0.004 | −0.009 | 0.001 | 0.185 |
| *GEO−BUT* | 0.007 | 0.002 | 0.012 | 0.001 |
| *BOM−BUT* | 0.061 | 0.056 | 0.067 | < 0.001 |
| *GEO−NOC* | 0.011 | 0.006 | 0.016 | < 0.001 |
| *BOM−NOC* | 0.065 | 0.059 | 0.071 | < 0.001 |
| *BOM−GEO* | 0.054 | 0.049 | 0.060 | < 0.001 |

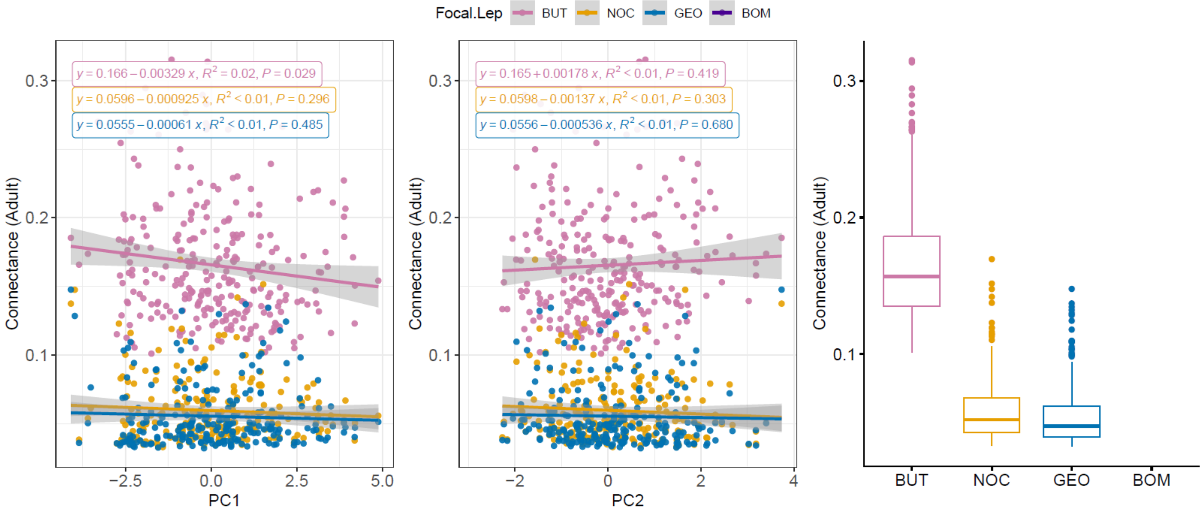


| | Df | Sum Sq | Mean Sq | F value | Pr(>F) |
|---|---|---|---|---|---|
| *Focal Lep.* | 2 | 2.011 | 1.006 | 998.896 | < 0.001 |
| *Residuals* | 743 | 0.748 | 0.001 | NA | NA |

| | Df | Sum Sq | Mean Sq | F value | Pr(>F) |
|---|---|---|---|---|---|
| *Focal Lep.* | 2 | 2.011 | 1.006 | 1007.576 | < 0.001 |
| *PC1* | 1 | 0.006 | 0.006 | 6.150 | 0.013 |
| *Focal Lep. × PC1* | 2 | 0.003 | 0.002 | 1.653 | 0.192 |
| *Residuals* | 740 | 0.738 | 0.001 | NA | NA |

| | Df | Sum Sq | Mean Sq | F value | Pr(>F) |
|---|---|---|---|---|---|
| *Focal Lep.* | 2 | 2.011 | 1.006 | 997.363 | < 0.001 |
| *PC2* | 1 | 0.000 | 0.000 | 0.002 | 0.967 |
| *Focal Lep. × PC2* | 2 | 0.002 | 0.001 | 0.929 | 0.395 |
| *Residuals* | 740 | 0.746 | 0.001 | NA | NA |

| | diff | lwr | upr | p adj |
|---|---|---|---|---|
| *NOC−BUT* | −0.106 | −0.112 | −0.099 | < 0.001 |
| *GEO−BUT* | −0.110 | −0.117 | −0.103 | < 0.001 |
| *GEO−NOC* | −0.004 | −0.011 | 0.003 | 0.329 |

Figure S12. GLM analyses on network connectance of the four focal clades, along PC1 (left panel) and PC2 (middle panel), or as a regional overall among-clade comparison (right panel), with life stages separated (upper: larva; lower: adult). See Fig. S8 for annotation explanations and detailed interpretation.

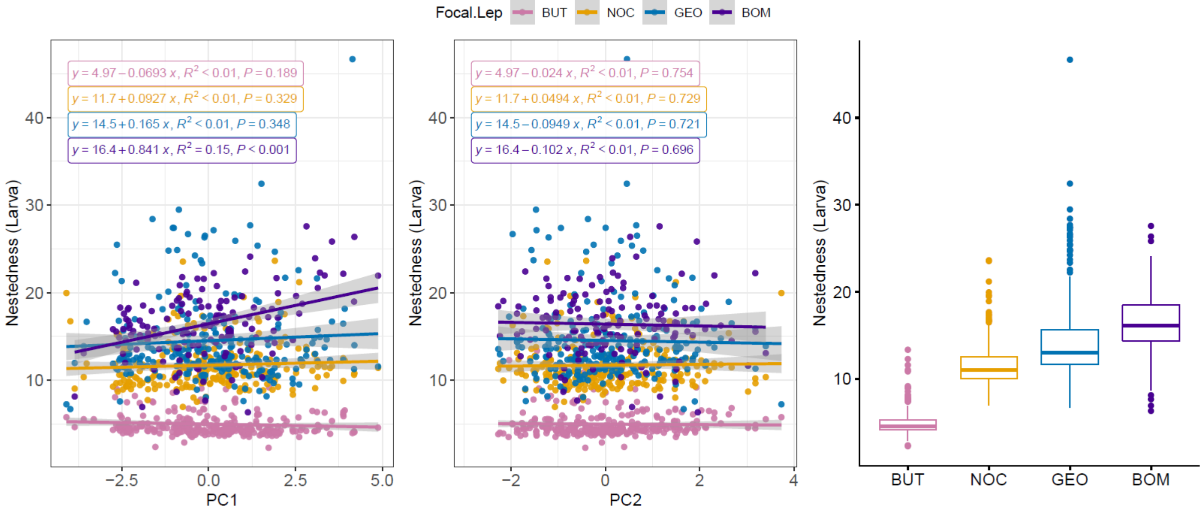


| | Df | Sum Sq | Mean Sq | F value | Pr(>F) |
|---|---|---|---|---|---|
| *Focal Lep.* | 3 | 17314.729 | 5771.576 | 529.662 | < 0.001 |
| *PC1* | 1 | 92.976 | 92.976 | 8.532 | 0.004 |
| *Focal Lep. × PC1* | 3 | 242.981 | 80.994 | 7.433 | < 0.001 |
| *Residuals* | 913 | 9948.705 | 10.897 | NA | NA |

| | Df | Sum Sq | Mean Sq | F value | Pr(>F) |
|---|---|---|---|---|---|
| *Focal Lep.* | 3 | 17314.729 | 5771.576 | 512.671 | < 0.001 |
| *PC2* | 1 | 1.655 | 1.655 | 0.147 | 0.701 |
| *Focal Lep. × PC2* | 3 | 4.584 | 1.528 | 0.136 | 0.939 |
| *Residuals* | 913 | 10278.423 | 11.258 | NA | NA |

| | Df | Sum Sq | Mean Sq | F value | Pr(>F) |
|---|---|---|---|---|---|
| *Focal Lep.* | 3 | 17314.73 | 5771.576 | 514.605 | < 0.001 |
| *Residuals* | 917 | 10284.66 | 11.216 | NA | NA |

| | diff | lwr | upr | p adj |
|---|---|---|---|---|
| *NOC−BUT* | 6.738 | 5.983 | 7.492 | < 0.001 |
| *GEO−BUT* | 9.546 | 8.792 | 10.299 | < 0.001 |
| *BOM−BUT* | 11.414 | 10.530 | 12.298 | < 0.001 |
| *GEO−NOC* | 2.808 | 2.039 | 3.576 | < 0.001 |
| *BOM−NOC* | 4.676 | 3.779 | 5.573 | < 0.001 |
| *BOM−GEO* | 1.868 | 0.972 | 2.765 | < 0.001 |

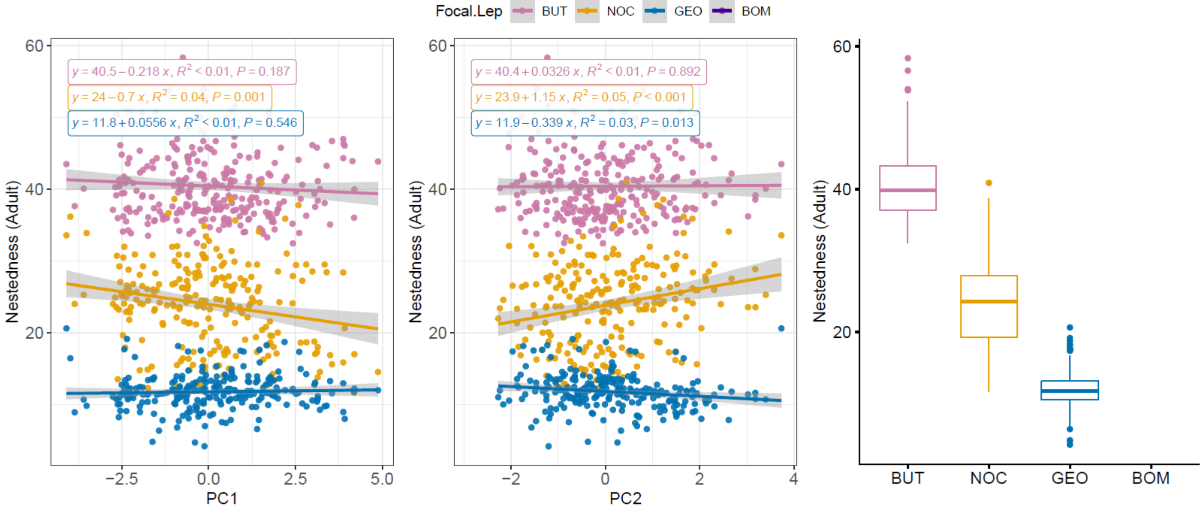


| | Df | Sum Sq | Mean Sq | F value | Pr(>F) |
|---|---|---|---|---|---|
| *Focal Lep.* | 2 | 104899.476 | 52449.738 | 2584.062 | < 0.001 |
| *PC1* | 1 | 184.435 | 184.435 | 9.087 | 0.003 |
| *Focal Lep. × PC1* | 2 | 214.362 | 107.181 | 5.281 | 0.005 |
| *Residuals* | 740 | 15020.078 | 20.297 | NA | NA |

| | Df | Sum Sq | Mean Sq | F value | Pr(>F) |
|---|---|---|---|---|---|
| *Focal Lep.* | 2 | 104899.476 | 52449.738 | 2595.866 | < 0.001 |
| *PC2* | 1 | 72.507 | 72.507 | 3.589 | 0.059 |
| *Focal Lep. × PC2* | 2 | 394.592 | 197.296 | 9.765 | < 0.001 |
| *Residuals* | 740 | 14951.776 | 20.205 | NA | NA |

| | Df | Sum Sq | Mean Sq | F value | Pr(>F) |
|---|---|---|---|---|---|
| *Focal Lep.* | 2 | 104899.48 | 52449.738 | 2527.432 | < 0.001 |
| *Residuals* | 743 | 15418.88 | 20.752 | NA | NA |

| | diff | lwr | upr | p adj |
|---|---|---|---|---|
| *NOC−BUT* | −16.431 | −17.380 | −15.482 | < 0.001 |
| *GEO−BUT* | −28.620 | −29.573 | −27.668 | < 0.001 |
| *GEO−NOC* | −12.189 | −13.171 | −11.208 | < 0.001 |

Figure S13. GLM analyses on network nestedness of the four focal clades, along PC1 (left panel) and PC2 (middle panel), or as a regional overall among-clade comparison (right panel), with life stages separated (upper: larva; lower: adult). See Fig. S8 for annotation explanations and detailed interpretation.

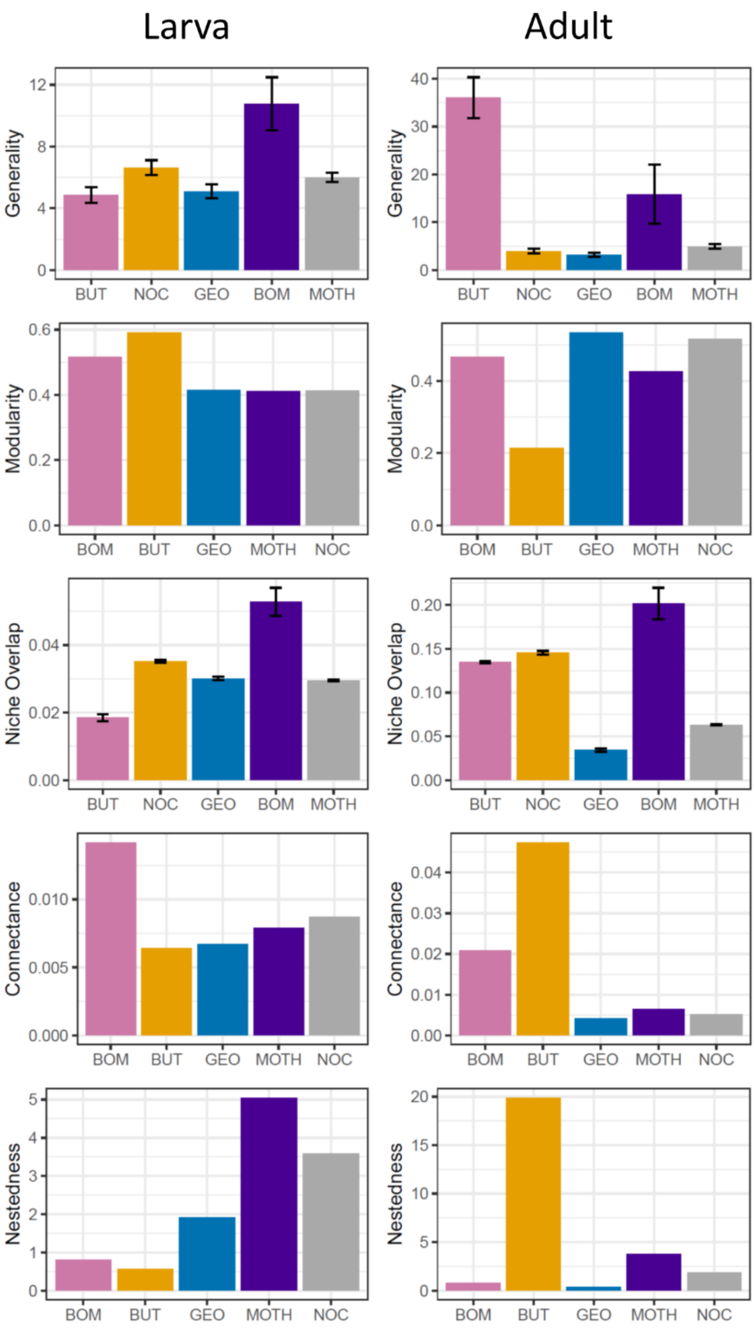


Figure S14. Comparisons of our selected network metrics among clades evaluated with the metawebs (subsets of respective Lepidoptera clade and life stage together the plants they interact with). Such comparisons consider only biological diets of Lepidopterans but not their realistic local co-occurring with food plants. Note that, for generality and niche overlap, the indices can be evaluated per Lepidoptera species, thus for the whole metaweb we can derive corresponding SE (error bars), while for the rest metrics there is only one evaluated value.

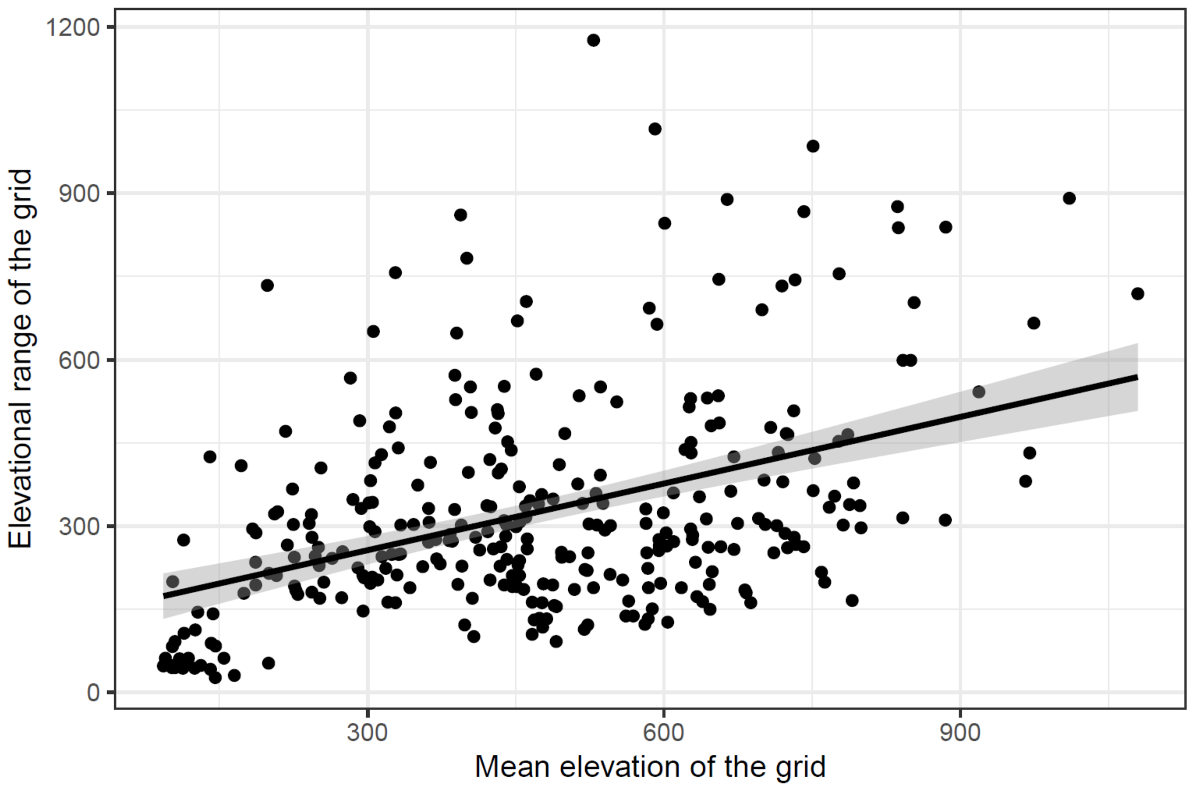


Figure S15. Across all the grids we studied, the elevational range per grid (i.e., the maximum minus the minimum elevation readings) increases with the mean elevation of the grid. This supports that topological variation within a grid can increase along with elevation.

**Section S3. References**


Aphalo P. (2022). ggpmisc: Miscellaneous Extensions to 'ggplot2'. R package version 0.4.7.

Auguie B. (2017). gridExtra: Miscellaneous Functions for "Grid" Graphics. R package version 2.3.

Dormann, C. F., Fruend, J., Bluethgen, N. & Gruber B. (2009). Indices, graphs and null models: analyzing bipartite ecological networks. The Open Ecology Journal, 2, 7-24.

Kassambara A. (2020). ggpubr: 'ggplot2' Based Publication Ready Plots. R package version 0.4.0.

Oksanen J., Simpson G., Blanchet F., Kindt R., Legendre P., Minchin P., O'Hara R., Solymos P., Stevens M., Szoecs E., Wagner H., Barbour M., Bedward M., Bolker B., Borcard D., Carvalho G., Chirico M., De Caceres M., Durand S., Evangelista H., FitzJohn R., Friendly M., Furneaux B., Hannigan G., Hill M., Lahti L., McGlinn D., Ouellette M., Ribeiro Cunha E., Smith T., Stier A., Ter Braak C., Weedon J. (2022). vegan: Community Ecology Package. R package version 2.6-2.

Schloerke B., Cook D., Larmarange J., Briatte F., Marbach M., Thoen E., Elberg A., Crowley J. (2021). GGally: Extension to 'ggplot2'. R package version 2.1.2.

Sievert, C. (2020). Interactive web-based data visualization with R, plotly, and shiny. CRC Press.

Thébault, E., & Fontaine, C. (2010). Stability of ecological communities and the architecture of mutualistic and trophic networks. Science, 329(5993), 853-856.

Urbanek S. (2013). png: Read and write PNG images. R package version 0.1-7.

Vu, Vincent Q. (2011) ggbiplot: A ggplot2 based biplot. R package version 0.55 755.

Wickham, H. (2016). ggplot2: Elegant Graphics for Data Analysis. Springer-Verlag New York.

Yu G. (2021). scatterpie: Scatter Pie Plot. R package version 0.1.7.
